# Refining type and timing of measured crop variables for the calibration of a new winter wheat cultivar in the STICS crop model

**DOI:** 10.1101/2025.02.10.637374

**Authors:** Meije Gawinowski, Maël Aubry, Samuel Buis, Cécile Garcia, Jean-Charles Deswarte, Marie-Odile Bancal, Marie Launay

## Abstract

Crop models need to be regularly upgraded with parametrization for new cultivars but this requires calibration, which is a major challenge. With winter wheat cultivar Rubisko as a case study, we propose to apply a calibration protocol to estimate the parameters of this new cultivar with multi-trials experimental data. We tested the calibration protocol in different conditions including or not LAI and/or biomass experimental data and we found that the resulting LAI and biomass dynamics strongly diverge. Several key findings emerge from this study: (1) RUE parameters should be excluded from the calibration process, as their critical role in biomass dynamics causes the optimization algorithm to treat them as adjustment parameters, resulting in unrealistic values for multiple parameters; (2) either LAI or biomass variables alone are sufficient for calibration, enabling experimental efforts to focus on one variable rather than both; and (3) the use of a synthetic dataset has facilitated the identification of the optimal type and timing of data collection needed to parameterize a new variety in the model. Moreover, the proposed methodology offers extrapolatable solutions applicable to other contexts (e.g., different models or datasets) and provides guidance on acquiring the most effective dataset for optimal calibration. The unbalanced structure of our dataset also highlighted the need to mobilize other calibration criteria (weighted RMSE) and alternative solutions to bridge the gap between quantitative metrics and empirical visual assessments.

## 1. Introduction

“Soil-crop” models are process-based models simulating crop growth, soil water and nitrogen dynamics in response to the environment and cultural practices. Developed since the 1960s, these models are widely used around the world for numerous purposes such as yield prediction, support to crop breeding and assessment/prediction of climate change impacts (Lobell and Asseng, 2017). Uncertainty in crop model simulations stems from model structure (i.e chosen formalisms of the different processes) and parameter values (Chapagain et al., 2022; Wallach and Thorburn, 2017). Some plant parameters can be generic while others are species- or cultivar-specific (Casadebaig et al., 2020). Regarding winter wheat, one of the major crops in the world and in France, phenology for example is highly heritable, well characterized by breeders, and usually firstly calibrated for new cultivars. Since crop models have to keep up with the evolution of cultural practices like choice of cultivar, it is necessary to regularly integrate new cultivars in the set of plant parameters. Doing so requires calibrating the model for these new cultivars, i.e. estimating model parameters on experimental data.

The aim of calibration is to estimate parameter values by minimizing the error between model outputs and observed data. This is a challenging exercise as many choices have to be made by the modeler which can impact the outputs, as highlighted in the framework of the AgMIP initiative (Agricultural Model Intercomparison Project; Rosenzweig et al., 2013) with the intercomparison of calibration practices among crop modelers (Wallach et al., 2021a):

– Choice of parameters to include and their range of variation, often based on modeler’s expertise and/or after sensitivity analysis
– Choice of target output variables, some of them might carry important experimental uncertainty
– Choice of the algorithm (e.g. frequentist vs Bayesian)
– Choice of the loss function
– Sequential vs all-together protocol
– Choice of calibration vs evaluation data sets

This intercomparison highlighted that these different choices led to heterogeneous calibration results, even for groups of modelers using the same model and same dataset. After this intercomparison, a common methodology was proposed and tested for phenology (Wallach et al., 2023) with a given simplex algorithm and given loss function for given target variables (phenological stages only) on observed French data. The modelers were allowed to choose the parameters to calibrate with a detailed documentation of the process, distinguishing major obligatory parameters from candidate parameters which are only included if their calibration reduces the BIC (Bayesian Information Criterion). The use of this common calibration methodology enabled to reduce variability between modelers but also prediction error. The next step aimed at generalizing this methodology to multiple types of data in crop models. To this end, a general methodology has recently been developed and tested on synthetic data by Wallach et al. (2024), who proposed a sequential approach for the calibration of different processes corresponding to different target variables and target parameters, which each step fed by the parameter values estimated in the previous step.

Parameter estimation is not straightforward as the number of parameters to optimize may be important whereas the amount of observations is limited, which is a common limitation when crop models are calibrated (Seidel et al., 2018). Many authors highlighted the importance of increasing the quantity and quality of the observed data from which depends the quality of the model calibration (Basso et al., 2010; Beaudoin et al., 2008; Dumont et al., 2014; Hernández-Ochoa et al., 2024). Moreover, the spatial and temporal patterns (especially in terms of soil, management and weather conditions) of the observations can be narrow, which may lead to larger errors when validating or applying the model as the calibration may not be representative in a wide range of environments (Hernández-Ochoa et al., 2024; Thorp et al., 2007). It is therefore appropriate to sparingly select the measurements to be collected so that they can be carried out in a wider range of environments. Research by Guillaume et al. (2011) specifically addressed this issue, demonstrating that dynamic variables should be included as target variables only if the goal is to enhance the model’s predictions for both dynamic and end-of-season variables, rather than solely for end-of-season variables. Since our goal is to calibrate the model for new wheat cultivars, and a significant part of the model’s application involves simulating carbon, nitrogen, and water dynamics, excluding dynamic variables like LAI and aboveground biomass is not possible. However, some of these dynamic variables are either time-consuming to assess, and so rarely available, or when assessed, covers a range of variables with different ecophysiological meanings that are more or less consistent with the simulated variables from the model. Leaf Area Index (LAI), which describes the amount of leaf area per unit of horizontal ground surface area, is a critical vegetation structural variable for modeling mass and energy exchange between the biosphere and the atmosphere (Yan et al., 2019). Fang et al. (2019) reviewed a large diversity of methods to measure LAI directly and indirectly. Direct measurements in the field consist mainly in destructive sampling of leaves, and have the disadvantage of being time-consuming and labor intensive which discourages large-scale sampling and long-term monitoring (Jonckheere et al., 2004). Indirect methods have therefore largely replaced them, using optical or remote sensing, and inferring LAI by measuring other variables such as gap fraction or light transmission (Tan et al., 2020; Weiss et al., 2004). They usually rely on the Beer-Lambert law, which describes the attenuation of light in uniform medium, and thus assimilate the canopy to an infinite turbid medium. This latter assumption is more or less consistent with crop canopies which can be highly structured such as maize or vineyards (Jiang et al., 2022), and the underestimation of LAI by Beer’s law in non-random distributed canopies is widely recognized (Yan et al., 2019). However, although these methods enable to monitor crops more effectively over space and time, they also introduce a high degree of uncertainty, as they include all the components of the canopy (stems, ears) and crop structure, which varies over time. Fang et al. (2019) also pointed out that the same term LAI could group together very different variables, such as GLAI (Green LAI), PAI (Plant Area Index, including stems and ears) or effective LAI (accounting for clumping effects), without this necessarily being defined; often parameterized on direct measurements, this introduces an additional uncertainty. Furthermore, Fang et al. (2019) summarized major effects that modify these indirect measurements at the time of measurement such as atmospheric conditions (diffuse to direct ratio, the height of the sun, radiation fluctuations, etc), background impact (soil reflectance), clumping (canopy structure), model inversion functions, and so on. As a result, major uncertainties arise from direct or indirect LAI measurements. Focusing on large-scale estimations, Fang et al. (2019) estimated to less than 1.37 m²/m² the RMSE in LAI assessed by moderate resolution remote sensing, correlating on average with reference LAI at r² =0.60. This performance was largely increased to RMSE < 0.83 m²/m² and r²=0.79 when the reference LAI was calibrated with high resolution remote sensing. For all these reasons, the calibration without LAI data should be tested.

In this paper, we thus propose to adapt the general method from Wallach et al. (2024) for the calibration of French cultivar Rubisko in the STICS crop model based on experimental data supplied by Arvalis. We propose to explore different calibration strategies in the framework of this method focusing on the availability of LAI and/or biomass observed data. We thus addressed two questions: 1) What set of parameters and target variables should be included in a calibration procedure by testing different combinations on experimental data and, 2) When experimental data should be acquired to build an efficient dataset for model calibration by testing different options first on synthetic data and then on experimental data.

## 2. Materials and Methods

### 2.1. Experimental data set

The target population was wheat fields in major wheat growing areas of France sown with winter wheat variety Rubisko and managed as usual. The crop measurement and management data as well as weather data were provided by ARVALIS - Institut du vegetal, a French agricultural technical institute, who run multi-year multi-purpose trials at multiple locations across France, which include variety trials. The trials follow standard agricultural practices. Weather data are from weather stations near each field and were completed by the SAFRAN meteorological reanalysis (Quintana-Seguí et al., 2008; Vidal et al., 2010) produced by the French national weather service (Meteo-France) when missing. The Rubisko data was separated into a calibration subset from 67 environments (17 sites, 7 years but not every year represented for every site) and an evaluation subset with data from 15 environments (13 sites, 4 years) by trying to maximize independence between the two subsets, but due to the near-complete design of the trials, there are some sites and years in common (but no site x year). The observed data were phenological dates for the beginning of elongation (BBCH30), flag leaf (BBCH 50), anthesis (BBCH 65) and maturity (BBCH 90), in-season LAI from sowing to anthesis, in-season aboveground plant biomass (t/ha) from sowing to harvest, end-of-season grain number (per m^2^) and grain yield (t/ha) at harvest.

### 2.2. STICS crop model and model initial conditions

STICS (V10.1.0) is a dynamic 1D soil-crop model that integrates crop development, growth, and yield formation with the carbon, nitrogen, energy, and water cycles of the soil-crop system (Beaudoin et al., 2023). It operates on a daily time-step using input data related to climate, crop species, soil conditions, agricultural management practices, and initial system states such as water and nitrogen content for each soil layer. Biomass growth is primarily driven by light interception, which is modeled using the big leaf approach based on leaf area index and the so-called Beer law of light extinction, combined with radiation-use efficiency. Crop development is influenced by thermal time, adjusted for vernalization and photoperiodic effects. Factors such as frost, nitrogen, suboptimal temperature or water deficiency can stress the crop, potentially impacting development, leaf area, growth and yield. Soil general characteristics (depth, water content at field capacity and wilting point, bulk density and other physical and chemical properties) were provided by Arvalis. The simulations start at the end of September (27th or 28th, i.e. day of year 270) a few weeks before sowing, the initial water availability is set at the field capacity on all horizons. Data from the white reports on cereals published in Belgium (https://livre-blanc-cereales.be/category/livreblanc/) and Beaudoin et al. (2005) are used to determine the initial nitrogen availability according to the previous crop.

### 2.3. Calibration method and protocol

To calibrate the STICS model, we employed the method proposed by Wallach et al. (2024), utilizing the R package CroptimizR (Buis et al., 2023) in combination with its wrapper function for STICS available in the R package SticsOnR (Lecharpentier et al., 2023) and the Nelder-Mead simplex algorithm that was used as in Wallach et al. (2024).

The proposed protocol by Wallach et al. (2024) is decomposed into preliminary steps (blue boxes in Figure 1) and a final step (red box in Figure 1). Each preliminary step corresponds to the calibration of a given process with a loss function based on ordinary least squares (OLS) and parameter selection with AICc. For each process, a set of major parameters is estimated. These parameters are chosen based on the assumption that they influence the simulated target-variables across nearly all environments. The parameter selection process is applied on an additional proposed set of candidate parameters that are supposed to explain a part of variability not explained by the major parameters. At each step, estimated parameters are then forced for calibration of the next process; for example, phenological parameters estimated during step 1 will be used for the next steps. Afterwards, the final step includes all the different processes and their target variables for the calibration of major and previously selected candidate parameters with a loss function based on weighted least squares (WLS) with initial parameter values set to previously estimated ones and weight values set to the error variances as estimated following the preliminary steps.

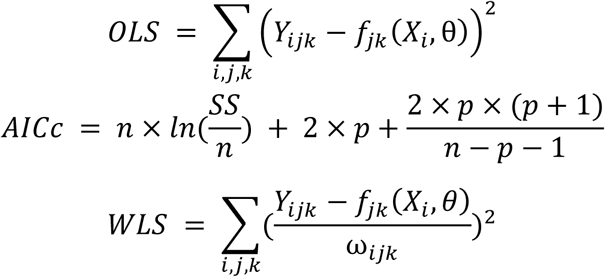

**Figure 1.**
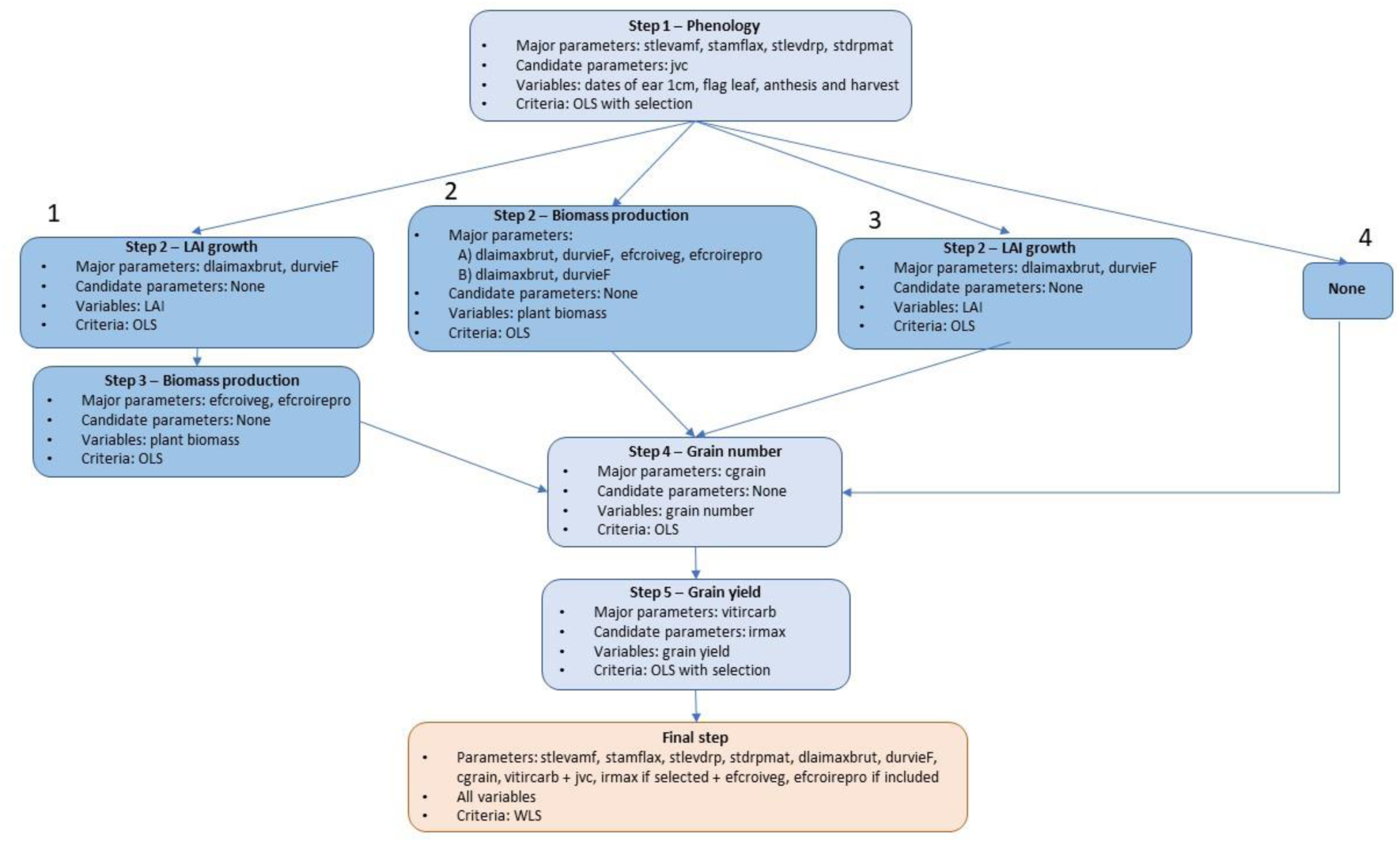
Flow diagram for the calibration protocol with different strategies for steps 2 and 3

where Y_ijk_ is the observed value for the k^th^ time point of the j^th^ variable in the i^th^ situation, f_jk_(X_i_; *θ*) the corresponding model prediction, ⍵_ijk_ is the weight and *θ* the vector of parameters to estimate, SS is the sum of squared errors for all variables in the group, n is the number of data points and p the number of estimated parameters.

The different steps are chosen depending on the availability of observed data and their order is based on their chronology and dependence level according to the model structure (chosen formalisms of the different processes), here phenology, LAI growth, production of plant biomass, grain number and grain yield at harvest. Since LAI and biomass data are not systematically measured in Arvalis trials, as their acquisition cost is higher, we explored different options for the second and third steps:

Strategy 1 - LAI and biomass data both available
Strategy 2 - Only biomass data available

A. Estimation of both RUE and LAI parameters
B. Estimation of LAI parameters only
Strategy 3 - Only LAI data available
Strategy 4 - No LAI nor biomass data available

The different calibration steps were chosen according to the work of Wallach et al. (2024). The choice of parameters to estimate for each step is based on the expertise of STICS modelers and their varietal aspect in the model. This choice is discussed in detail in the discussion subsection 4.1. The default values for these parameters correspond to the value of the already calibrated cultivar Arminda, which is analogous to Rubisko in terms of precocity. The steps with their dedicated target variables and parameters to estimate are detailed in Figure 1. The list of major and candidate parameters as well as their definition, units, bounds and default values are detailed in Table 1.

**Table 1.**
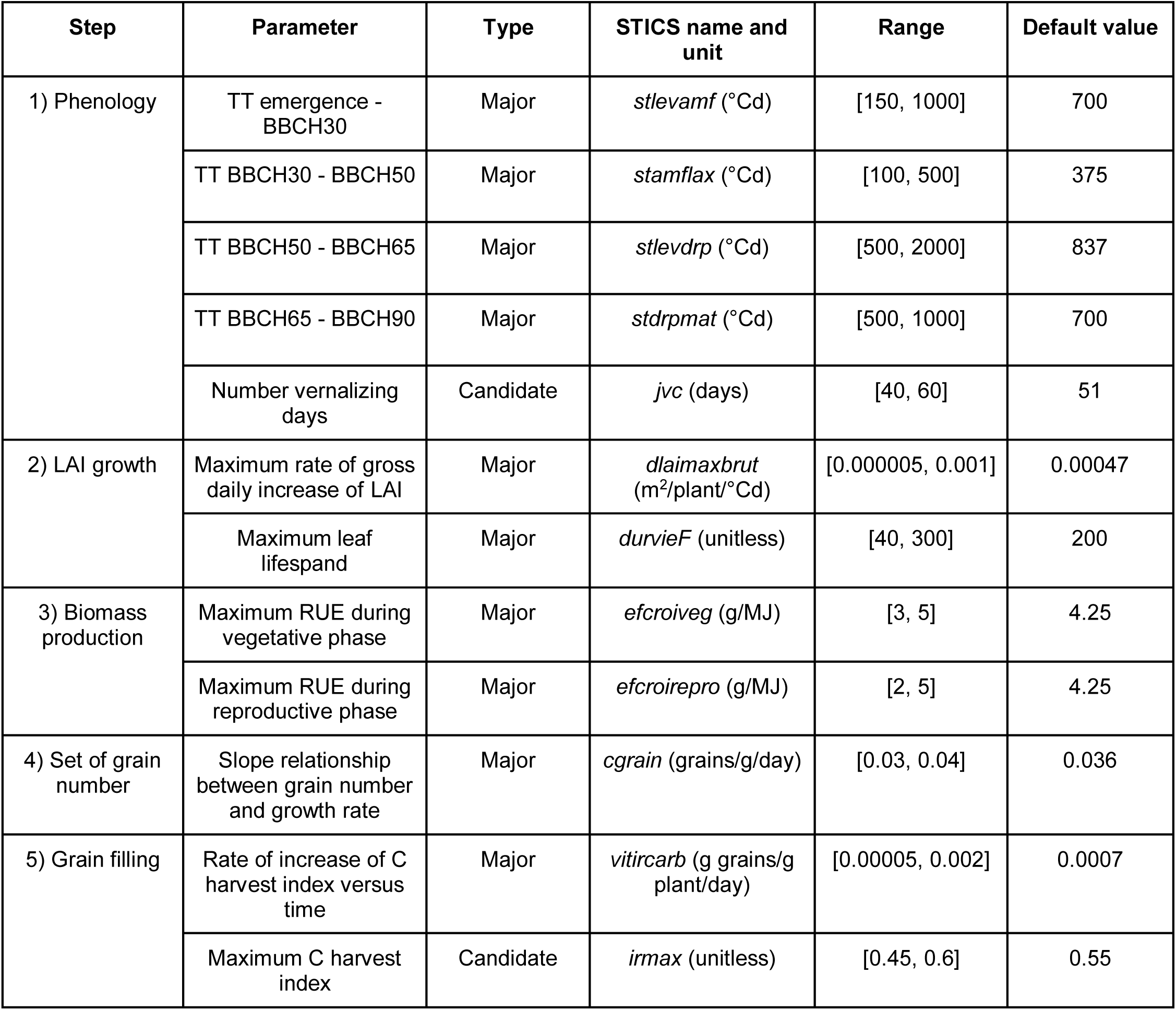
STICS parameters selected for calibration.

| Step | Parameter | Type | STICS name and unit | Range | Default value |
| --- | --- | --- | --- | --- | --- |
| 1) Phenology | TT emergence - BBCH30 | Major | *stlevamf* (°Cd) | [150, 1000] | 700 |
|  | TT BBCH30 - BBCH50 | Major | *stamflax* (°Cd) | [100, 500] | 375 |
|  | TT BBCH50 - BBCH65 | Major | *stlevdrp* (°Cd) | [500, 2000] | 837 |
|  | TT BBCH65 - BBCH90 | Major | *stdrpmat* (°Cd) | [500, 1000] | 700 |
|  | Number vernalizing days | Candidate | *jvc* (days) | [40, 60] | 51 |
| 2) LAI growth | Maximum rate of gross daily increase of LAI | Major | *dlaimaxbrut* (m<sup>2</sup>/plant/°Cd) | [0.000005, 0.001] | 0.00047 |
|  | Maximum leaf lifespan | Major | *durvieF* (unitless) | [40, 300] | 200 |
| 3) Biomass production | Maximum RUE during vegetative phase | Major | *efcroiveg* (g/MJ) | [3, 5] | 4.25 |
|  | Maximum RUE during reproductive phase | Major | *efcroirepro* (g/MJ) | [2, 5] | 4.25 |
| 4) Set of grain number | Slope relationship between grain number and growth rate | Major | *cgrain* (grains/g/day) | [0.03, 0.04] | 0.036 |
| 5) Grain filling | Rate of increase of C harvest index versus time | Major | *vitircarb* (g grains/g plant/day) | [0.00005, 0.002] | 0.0007 |
|  | Maximum C harvest index | Candidate | *irmax* (unitless) | [0.45, 0.6] | 0.55 |

The performance of these calibration strategies is assessed by:

– Comparison of estimated values for the different parameters
– Visual comparison of simulated vs observed variables
– Computation of normalized RMSE for each variable v (NRMSEv) and globally (NRMSE) for each environment i such as done in Guillaume et al. (2011)with a comparison of their distributions and means:

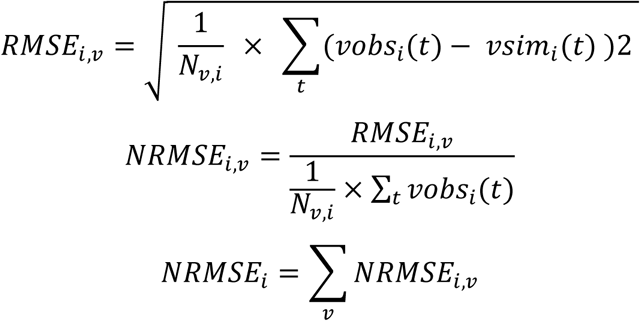
– Computation of correlation indices for the different variables between observed and simulated variables
– Unlike NRMSE, MSE can be decomposed into bias and difference in variability as in Guillaume et al. (2011):

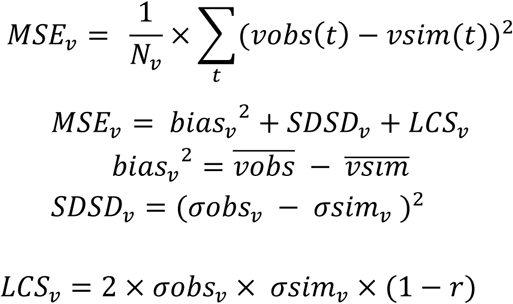

where 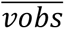 is the mean simulated values, *σobs* and *σsim* are the standard deviations of the observed and simulated values and r is the correlation between observations and simulations. The bias component expresses the average difference between observations and simulations, the SDSD component expresses the difference in variability between observations and simulations and the LCS component is the residual with no interpretation. Here the decomposed MSE is computed for each variable for the entire dataset so vobs (respectively vsim) contains the observations (resp. simulations) of all the environments, whether for the computation of individual NRMSEs vobs (respectively vsim) only contains the observation (resp. simulations) for environment i.

### 2.4. Effect of data availability using synthetic data for LAI or biomass

The calibration performance is not only related to the calibration protocol but also to data availability, i.e. the number of observed variables as well as the number and repartition of observations throughout growth for each variable. The dataset provided by Arvalis for the test of different calibration strategies enabled to perform calibration including LAI and/or biomass data, however the repartition of observations is heterogeneous for both variables: LAI observations are concentrated during LAI growth but there are only a few points around maximum LAI and none during LAI senescence, for biomass there are very few points at the end of growth around maturity. As this heterogeneity in the timing of data acquisition can impact calibration performance, we propose to explore the effects of different temporal data repartition for LAI or biomass separately on calibration performance using synthetic data. The question studied here is: what temporal distribution of LAI and biomass measurements yields the best calibration results? Here LAI and biomass are studied separately to represent the situation where only LAI or only biomass data are available.

Plant growth is divided in three stages based on LAI dynamics: before the peak (35 to 105 day of year), at peak (110 to 155 day of year), after the peak (160 to 300 day of year) and throughout growth (Figure 2). Synthetic data were generated using the parameter values estimated from the calibration on the real data. Five values for LAI or biomass were sampled at regular time intervals within the time slot considered for each one the 67 environments of the calibration dataset and 15 environments of the validation dataset. A Gaussian random noise of mean 0 and standard deviation equal to 10% of mean simulated values were added to the simulated values. Calibration strategies depend on data availability but also the conclusion from the testing on real observed data; for LAI synthetic data was simulated with parameters values estimated using the “LAI” strategy (Fig.1, strategy 3) whereas synthetic biomass data was simulated with parameters values estimated using the “Biomass (no RUE)” strategy (Fig.1, strategy 2B). In both cases, real data from the original calibration dataset were added to the synthetic dataset for the other variables (phenology, grain number and grain weight). As for the previous section, initial parameter values are set to default values of cultivar Arminda.

**Figure 2.**
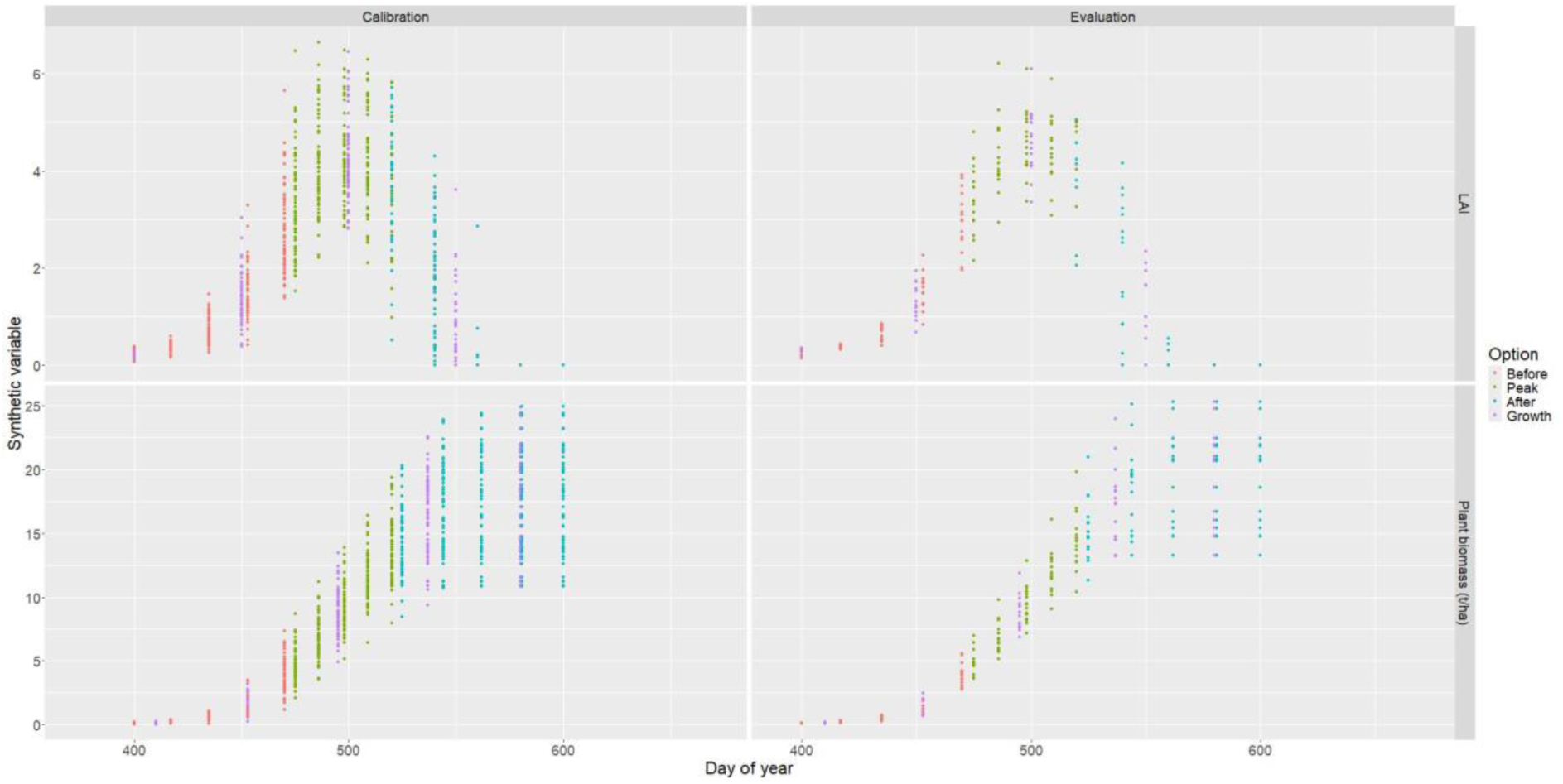
Synthetic data for LAI and biomass with different options of data availability (before, during or after the LAI peak, or throughout growth) for the 67 environments of cultivar Rubisko.

The results obtained for the four options of LAI and biomass data availability (data available “Before”, during “Peak”, “After” peak or throughout “Growth”) will be compared for LAI and biomass based on NRMSE values as in Guillaume et al. (2011).

### 2.5. Calibration on a subset of observed LAI or biomass data

After exploring different options for LAI and biomass data availability on synthetic data (see 2.4 for description), the best option (“Peak”, see results 3.2) was selected to subset actual observed data and compare calibration performance between entire and subsetted data. For synthetic exploration, peak data was defined by a temporal window (between 110 and 155 day of year), however, as there are very few LAI points in this temporal window for the actual observed data (only 6 for 4 environments), observed LAI data at peak was rather sampled above a threshold value of 2.5. Biomass points were sampled in the peak temporal window as for the synthetic exploration since there are sufficient observations, with a maximum of 5 points per environment. As for the synthetic exploration, calibration is performed with the “LAI” strategy for LAI data and “Biomass (no RUE)” for the biomass data.

## 3. Results

### 3.1. Comparison of different calibration strategies on observed experimental data

#### 3.1.1. Values of estimated parameters

Phenological parameters (*stlevamf*, *stamflax*, *stlevdrp*, *stdrpmat* and *jvc*) are similarly estimated across the different strategies (Figure 3). LAI parameters (*dlaimaxbrut*, *durvieF*) are estimated at contrasted values between the different strategies, leaf lifespan *durvieF* is notably estimated to the upper bound at 300°Cd for the strategy “Biomass (no RUE)”. For biomass parameters, i.e. radiation-use-efficiency parameters (vegetative RUE *efcroiveg*, et reproductive RUE *efcroirepro*), when calibrating only on biomass data (strategy “Biomass (with RUE)”), *efcroiveg* is estimated at the upper bound at 5 g/MJ which is not considered an attainable value in terms of physiology for wheat species. Besides, with this strategy LAI parameters are estimated at low values resulting in an underestimated dynamic based on visual assessment (Figure 4). We hypothesize that when calibrating LAI and RUE parameters on biomass data, the algorithm favors adjusting RUE as the easiest (i.e. most direct) way to impact biomass. However reproductive RUE *efcroirepro* is estimated at quite low values, resulting in biomass dynamics visually underestimated. For grain number, the *cgrain* parameter is calibrated with little variation between the different calibration strategies. For grain weight, the estimated values for *irmax* show little variation between the different calibration strategies whereas there are different estimated values for *vitircarb*, demonstrating that this parameter compensates for different levels of biomass production among the different calibration strategies.

**Figure 3.**
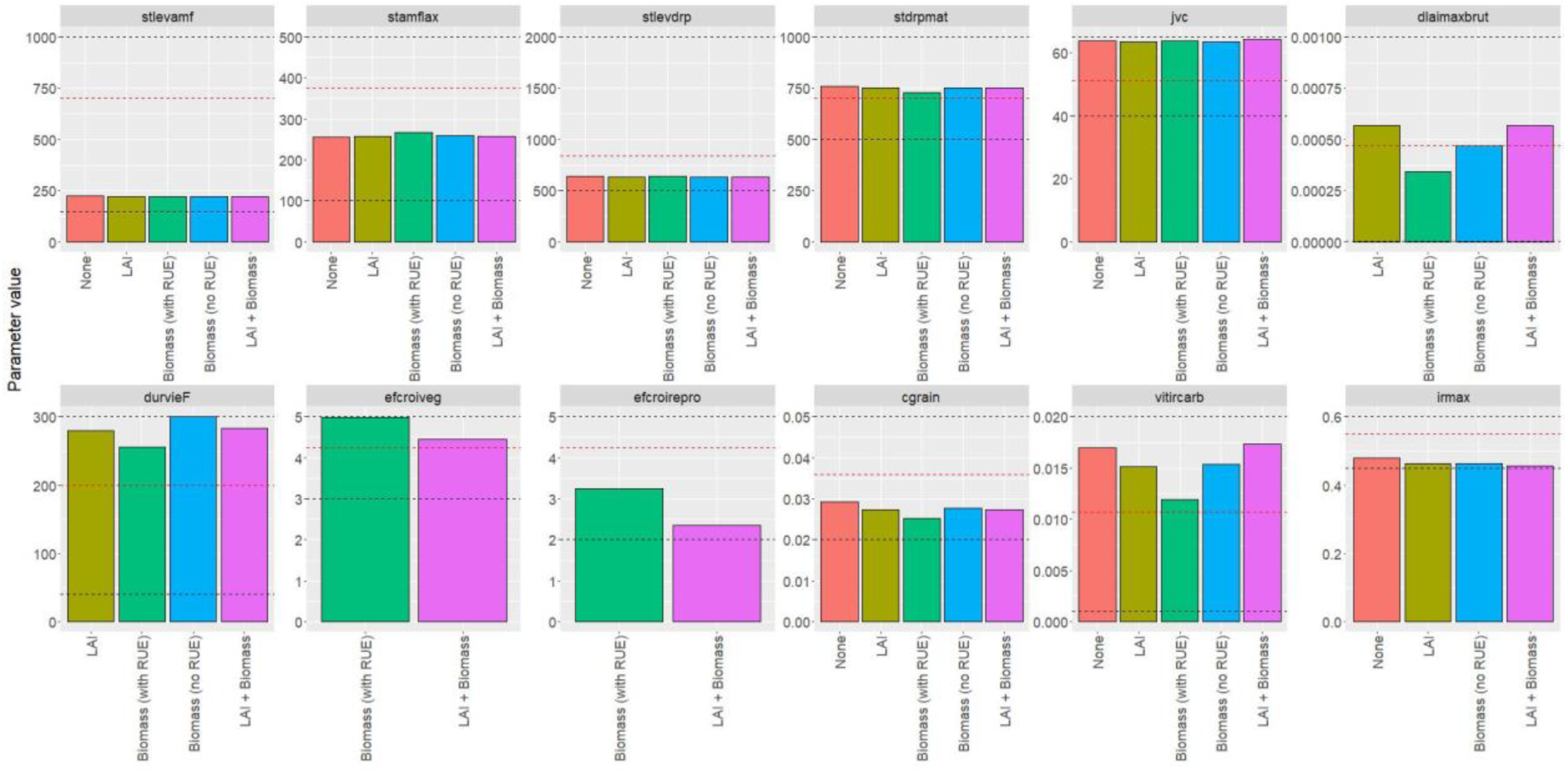
Estimated values of calibrated parameters for each calibration strategy. Black dashed lines represent lower and upper bounds, red dashed lines represent default values.

**Figure 4.**
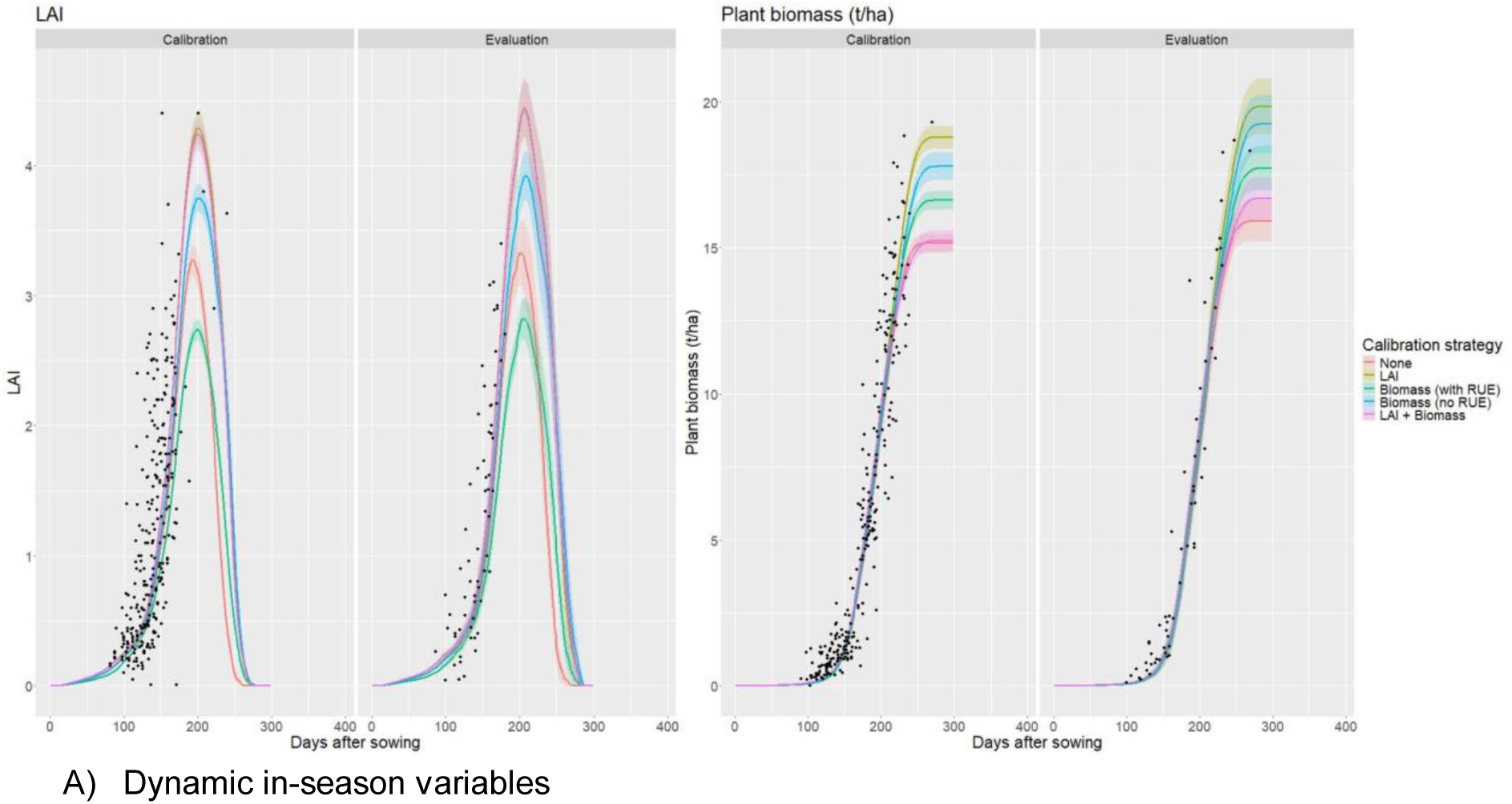
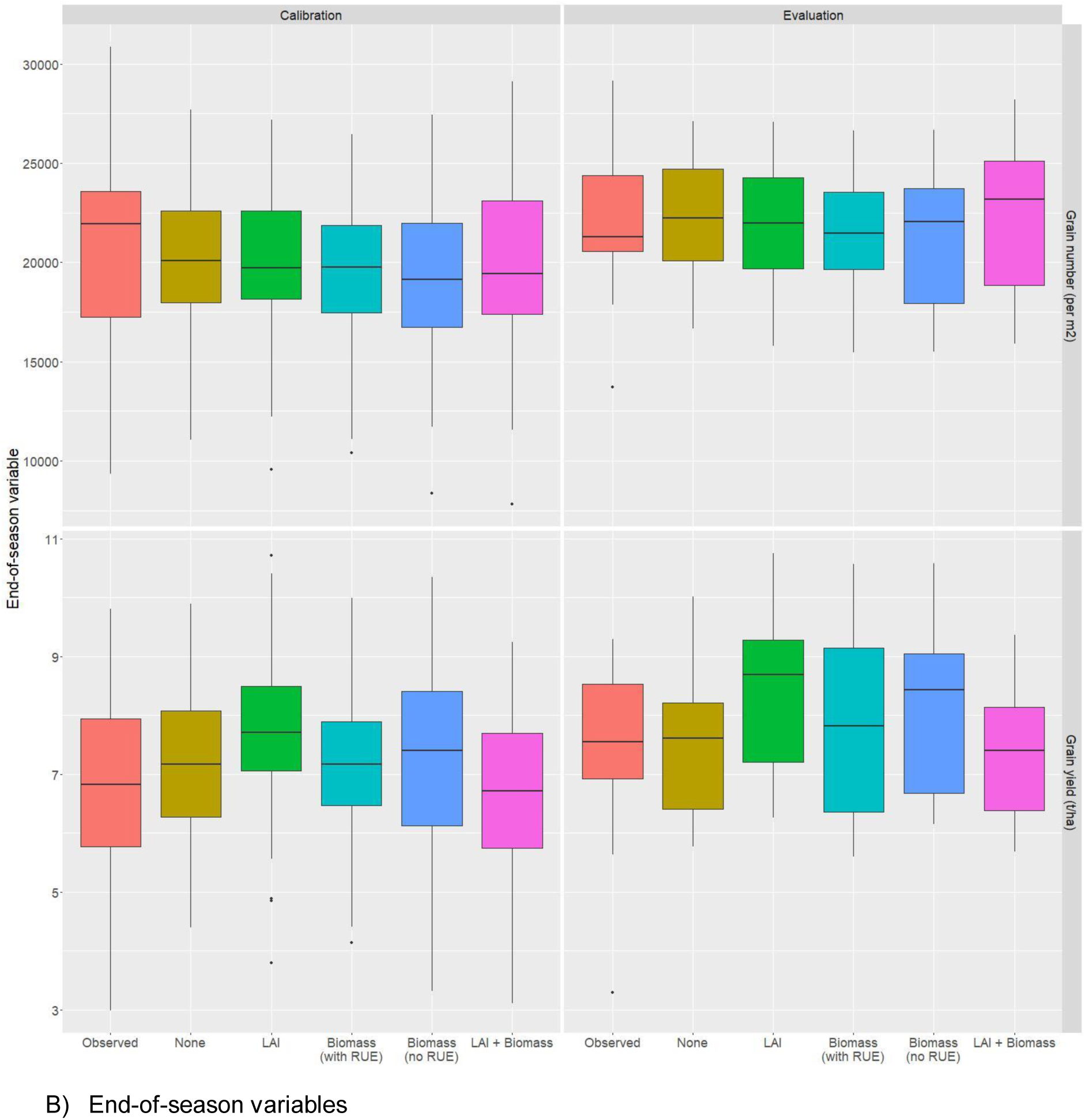
Observed and dynamic variables observed and simulated by STICS for the calibration and evaluation data sets A) in-season and B) end-of-season. For in-season variables (A) black points represent observations, coloured lines represent the average of the variables over the different simulations for each calibration strategy and the colored ribbon represents the standard error around the mean. The distribution of end of seasons variables (B) is shown as box plots showing median, quartiles and dispersion, in the case of observed values (red) and the different strategies of calibration, that is none (olive green), LAI (green), biomass with RUE (green-blue), biomass without RUE(marine blue) and LAI+Biomass (pink)

#### 3.1.2. Visualization of goodness-of-fit

Fits of simulations with parameters estimated with the different calibration strategies are displayed on Figure 4. As expected, LAI simulations are better fitted when the model is calibrated with LAI data (on observed LAI alone or observed LAI + biomass). However, there are very few LAI observed points at maximum LAI so it is difficult to visually assess the goodness-of-fit of the simulated LAI with the observed maximum LAI, and at this stage the “Biomass (no RUE)” strategy also gives good results (Figure 4A). In contrast, the “Biomass strategy (with RUE)” clearly underestimates the LAI, which is directly related to the underestimation of LAI parameters with this strategy as shown above. Moreover, the late biomass simulations are underestimated for the “Biomass (with RUE)” and “LAI+Biomass” strategies, which is directly related to the underestimation of the reproductive RUE parameter *efcroirepro* under these strategies. On the other hand, the “LAI” strategy (on the calibration and evaluation datasets) and the “Biomass (no RUE)” strategy (on the evaluation dataset) seemed to produce the most realistic simulations of dynamic biomass. Also, biomass simulations are very similar with low variability until about 225 days after sowing but diverge substantially at maturity with a difference of nearly 4 t/ha (Figure 4A). The residual plot (Figure S1) confirms the better performance of the “LAI” and “Biomass (no RUE)” strategies in reproducing the dynamics of LAI and plant biomass at the end of the season, as well as the scarcity of observed data at late crop growth and during maturity. We also note that above the LAI value of 2.5, the divergence in performance between strategies (with relatively stable residuals for the “LAI”, “LAI+Biomass” and “Biomass (no RUE)” strategies and increasing residuals for the “Biomass (with RUE)” and “None” strategies) is associated with the drastic reduction in the number of observations. This lack of crucial data could explain why the visualization of goodness of fit is so delicate and sometimes different between calibration and evaluation datasets.

Observed yield and grain number are highly variable, so this is difficult to visually assess goodness-of-fit (Figure 4B) but the “None” and “LAI + Biomass” strategies show similar medians and close dispersions, whereas they are overestimated with the other strategies. However, since there can be compensation through the estimation of yield and grain number parameters, the evaluation of these variables is less crucial than plant biomass or LAI. Phenology is similarly estimated with the different calibration strategies with a satisfactory goodness-of-fit (Figure S2).

#### 3.1.3. Comparison of NRMSE

In addition to visual assessment, individual NRMSE values were also computed to compare the different calibration strategies for LAI, plant biomass, grain number, grain yield and all the output variables together: their means for each group (strategy x dataset type x variable) can be found in Table 2 for both the calibration and evaluation datasets. Their distributions are also represented on Figure S3.

**Table 2.** Mean individual NRMSE values for LAI, biomass, grain number, grain weight and all variables for the calibration and evaluation datasets for each calibration option. Bold values are the minimal ones for calibration and evaluation.

| Calibration option | Dataset type | Output variable |  |  |  |  |
| --- | --- | --- | --- | --- | --- | --- |
|  |  | LAI | Biomass | Grain number | Grain weight | All |
| None | Calibration | 0.50 | **0.22** | 0.15 | **0.18** | **1.08** |
|  | Evaluation | 0.55 | **0.24** | **0.13** | **0.15** | **1.11** |
| LAI | Calibration | **0.48** | **0.22** | 0.16 | 0.22 | 1.12 |
|  | Evaluation | 0.50 | **0.24** | 0.14 | 0.25 | 1.16 |
| Biomass with RUE | Calibration | 0.55 | **0.22** | **0.14** | 0.19 | 1.13 |
|  | Evaluation | 0.63 | **0.24** | **0.13** | 0.20 | 1.24 |
| Biomass no RUE | Calibration | 0.49 | 0.23 | 0.18 | 0.20 | 1.13 |
|  | Evaluation | 0.53 | **0.24** | 0.15 | 0.22 | 1.17 |
| LAI + Biomass | Calibration | **0.48** | 0.24 | 0.19 | 0.21 | 1.15 |
|  | Evaluation | **0.49** | **0.24** | **0.13** | **0.15** | **1.11** |

The distributions of individual NRMSE values for biomass are very similar between the different strategies with similar mean values (Table 2) but they diverge for LAI, grain number, grain yield and global NRMSEs. Indeed LAI shows a lower mean NRMSE for the strategies including LAI data in the calibration process (“LAI”, “LAI + Biomass”). For grain number, grain yield and all variables, mean NRMSEs are lower for the strategies “None” and “LAI + Biomass”. Correlations between observed and simulated data were also computed (Table S1), with very similar results to NRMSE values.

For LAI, visual fit is consistent with individual NRMSE values, leading to the conclusion that LAI is better predicted for calibration strategies including LAI data. However, for plant biomass, the analysis of individual NRMSE values differs from visual analysis; individual NRMSE values for plant biomass are very similar whereas visually some calibration strategies (“None”,“LAI + Biomass” and “Biomass (with RUE)” to some extent) clearly seem to simulate underestimated plant biomass (Figure 4A).

#### 3.1.4. Decomposition of MSE into bias and difference in variability

MSE was then decomposed into bias and difference in variability (section 2.3.) and this decomposition is represented on Figure 5. Note that MSE is computed on the calibration and evaluation dataset as a global error and not as the mean of individual errors as for the previous section, so conclusions can differ. For the calibration dataset, the bias for LAI is much higher for the “Biomass (with RUE)” strategy as LAI is strongly underestimated (Figure 4) and then simulated biomass is further compensated with overestimated RUE value (Figure 3); on the contrary this bias is reduced for strategies including LAI data. For plant biomass MSE values are more variable than the mean of individual NRMSEs, with lower values for the strategies “Biomass (with RUE)” and “LAI + Biomass”, which also contradicts visual analysis. For grain yield, the “LAI” strategy shows a higher MSE value due to much higher bias. For the validation dataset MSEs are similar to calibration for grain yield but higher than for calibration, due to a higher bias for both LAI and plant biomass. However, for grain number, MSEs are smaller and less variable across strategies than for the calibration dataset.

**Figure 5.**
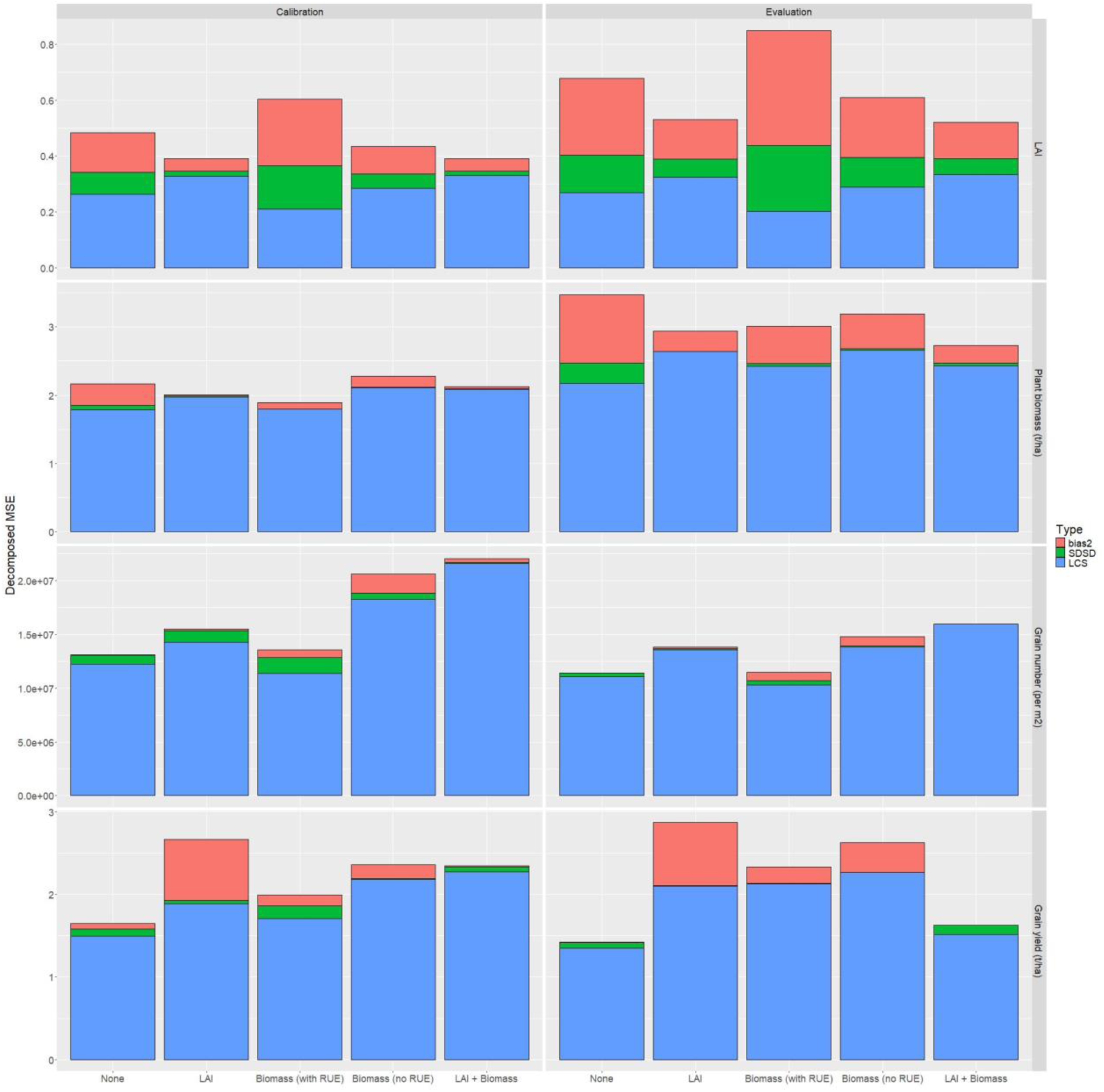
Decomposed MSE into bias (bias2, in red), difference in variability (SDSD, in green) and residual (LCS, in blue) faceted for LAI, plant biomass, grain number and grain yield for the different calibration strategies computed on the calibration and evaluation datasets.

Among the different calibration strategies tested in this section, none is perfect. We identified that:

– the “biomass (with RUE)” strategy resulted in a large underestimation of LAI parameters and an overestimation of the vegetative RUE parameter, as well as an underestimation of plant biomass dynamics. This strategy should therefore be abandoned;
– the “LAI + Biomass” strategy, although interesting in terms of NRMSE and decomposed MSE to simulate LAI, grain number and yield (more specifically on the evaluation dataset), underestimates biomass at the end of the season;
– the “LAI” and “Biomass (no RUE)” strategies emerge as the best options for this dataset; they provide good dynamics of LAI and plant biomass, and both satisfy NRMSE and reasonable bias in the decomposed MSE for the calibration and evaluation datasets (except for grain yield, which highlights a high bias with the “LAI” strategy).

### 3.2. Comparison of different options of temporal availability for LAI or biomass data

#### 3.2.1. Comparison of calibration performance in response to data availability on synthetic data

In this section we study the temporal availability of LAI or biomass data separately. For LAI, the only possible calibration strategy is the “LAI” one. For plant biomass the “Biomass (no RUE)” strategy was finally agreed upon in the previous subsection on observed data. Hence, for LAI (respectively plant biomass), the “LAI strategy (resp. “Biomass (no RUE)”) strategy is used to simulate synthetic data and then perform calibration.

Simulations performed with estimated parameters for the different options for data availability are very similar, only few differences can be distinguished around maximum LAI for LAI simulations, and at maturity for biomass simulations (Figure 6), as well as grain number and grain weight (data not shown). This reflects the very similar estimated parameter values among the different options (Figure S4), even though there are some differences on different types of parameters (phenology, LAI, yield).

**Figure 6.**
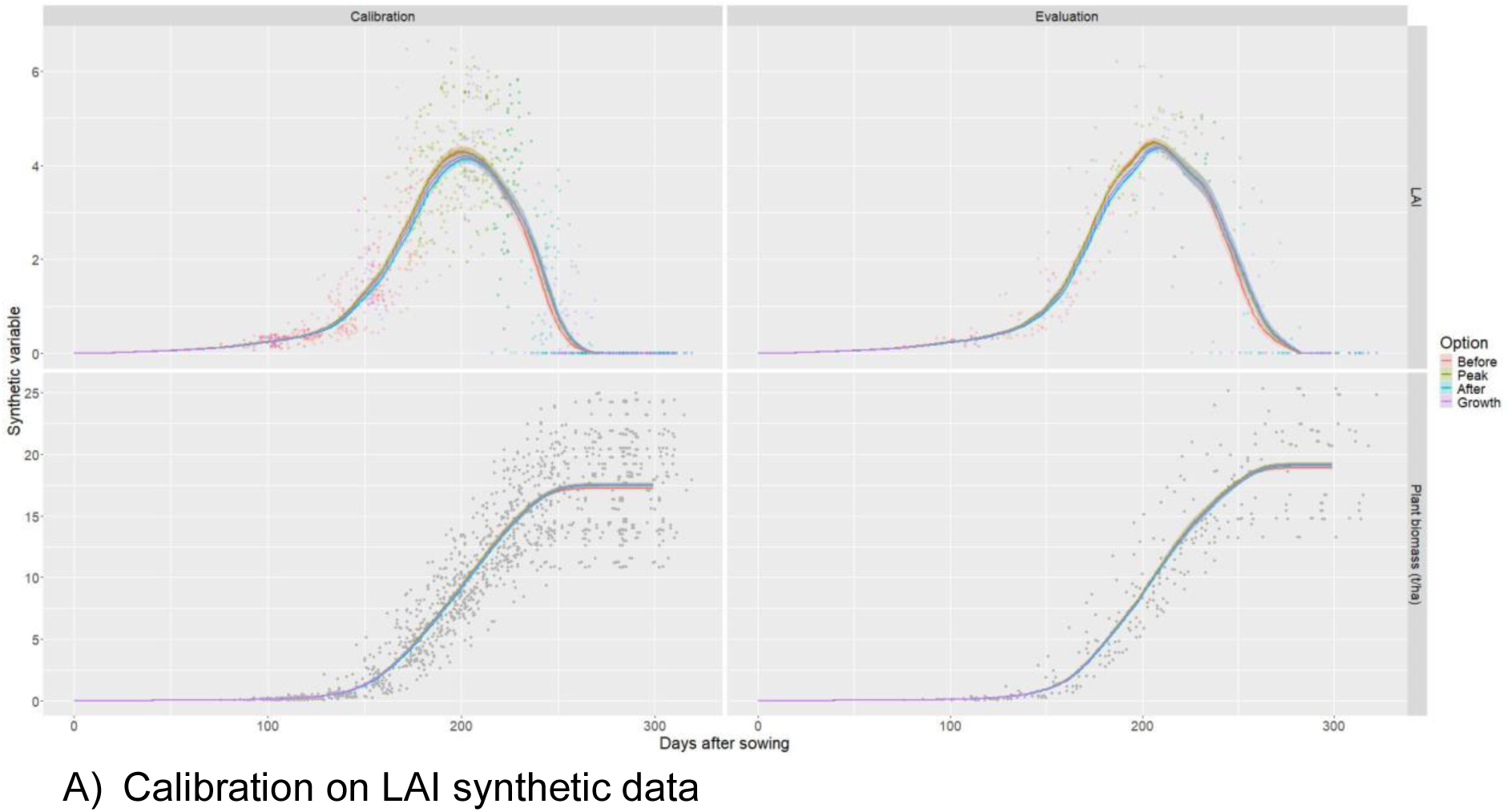
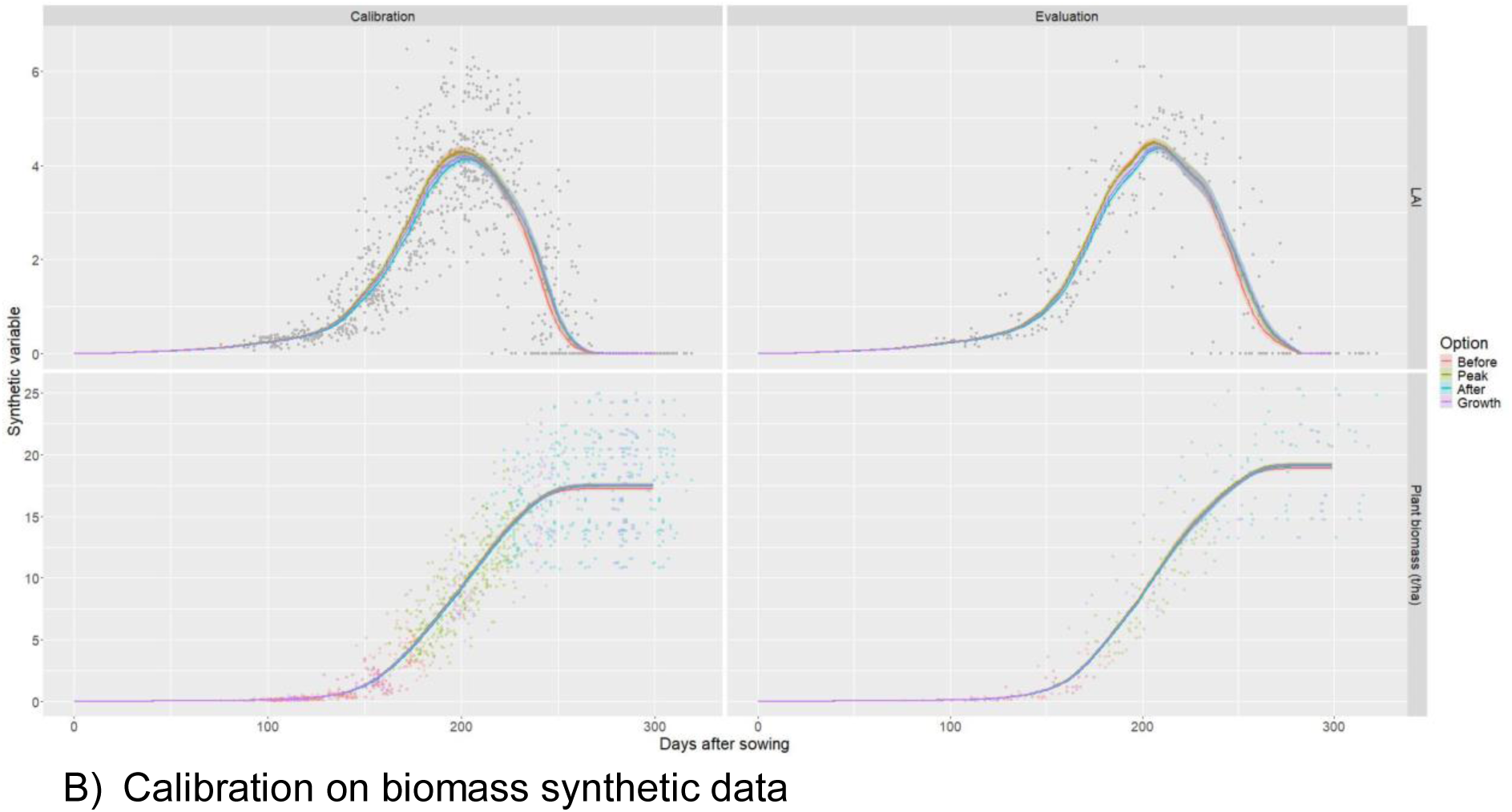
Simulations (continuous lines) with parameters estimated with different strategies for data availability (colors) and observations (points) for LAI and plant biomass. Grey points indicate observations not used in the calibration.

The distributions of NRMSE values are relatively similar for LAI and the magnitude of the errors remains low (Figure 7). However, it is worth noting that the median of the NRMSE is lower for the Peak strategy. For biomass there is a clear distinction between the “Before” strategy with considerably higher NRMSE values, whereas the “Peak” strategy displays low values with low variability, even if again, the magnitude of errors is very low (Figure 7).

**Figure 7.**
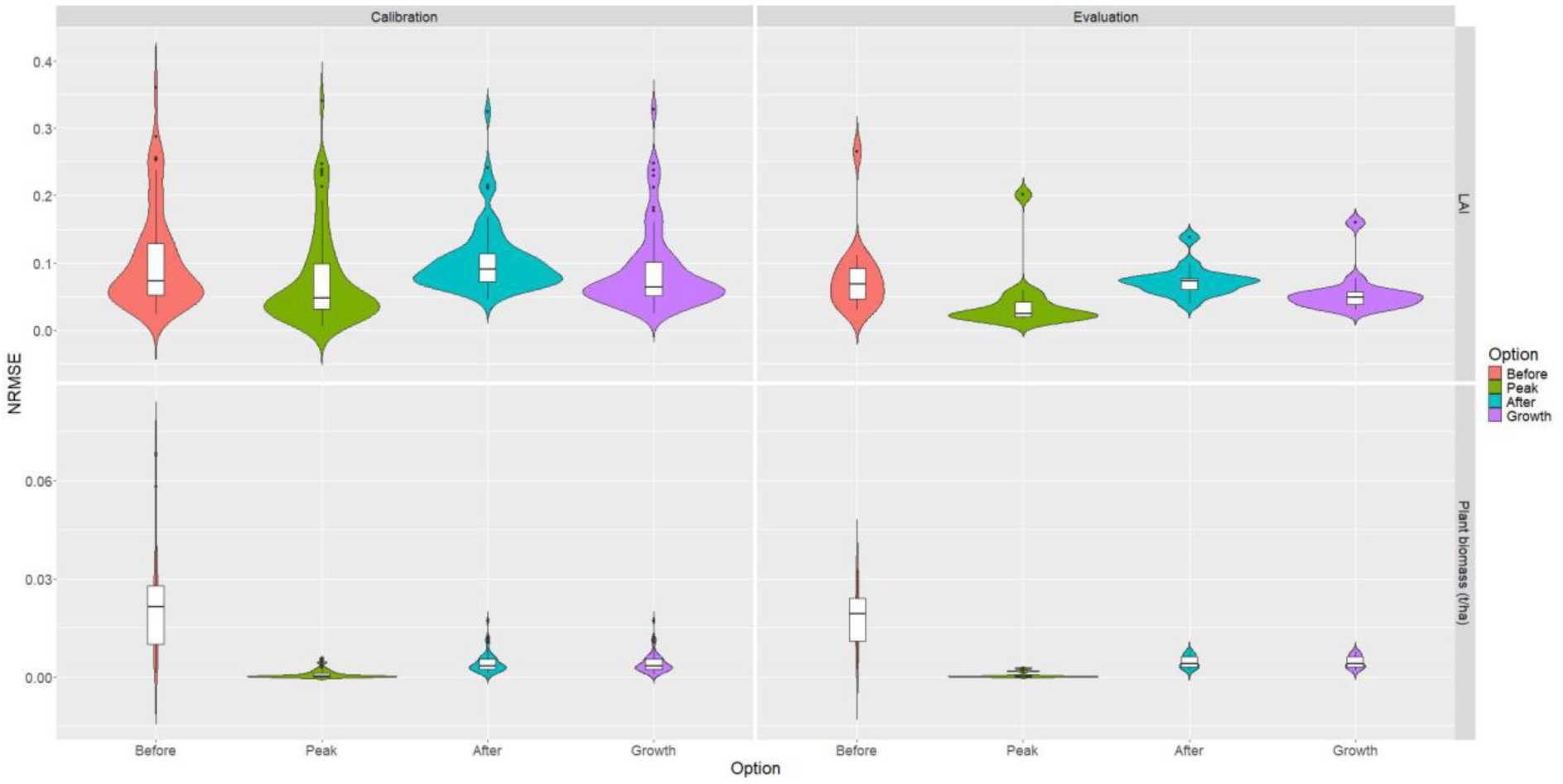
Distribution of NRMSE values for LAI and plant biomass between different calibration strategies for data availability.

#### 3.2.2. Calibration with observed LAI or biomass data at peak

In the previous section we found that calibrating on LAI or biomass synthetic data around maximum LAI was the best option with lower NRMSEs. As a result, this sampling around maximum LAI was then applied to the observed data for LAI and plant biomass separately so that the results can be compared with the ones obtained with synthetic data. Results obtained after a calibration performed on the entire observed data are hereafter referred to as the “All” option whereas results obtained with data sampled around the peak are referred to as the “Peak” option.

We can observe on Figure 8 that simulated LAI and biomass dynamics are lower when calibrated on peak data (“Peak” option) compared to the entire data (“All” option), resulting in a visual underestimation. These trends are due to lower estimated values for the leaf lifespan duration parameter *durvieF* (Figure S5). When comparing NRMSE values (Table 3), they are indeed lower for the “Peak” option when computed on the peak data, but they are higher for the “Peak” option when computed on the entire data (which is the most expected performance). As expected, mean NRMSE values computed on the peak data are indeed lower with the “Peak” option. Mean biomass NRMSE values computed on the entire data are also lower with the peak option but it is not the case for LAI. Even though NRMSE values are lower with the “Peak” option, the resulting simulated LAI peak (Figure 8A) and end-of-season biomass appear underestimated. This contradiction between visual analysis and NRMSE values is probably due to the low number of observations for these periods of time, but also to the difference in data structure between observed and synthetic data (balanced design between the environments resulting in a higher number of observations sampled at regular intervals).

**Figure 8.**
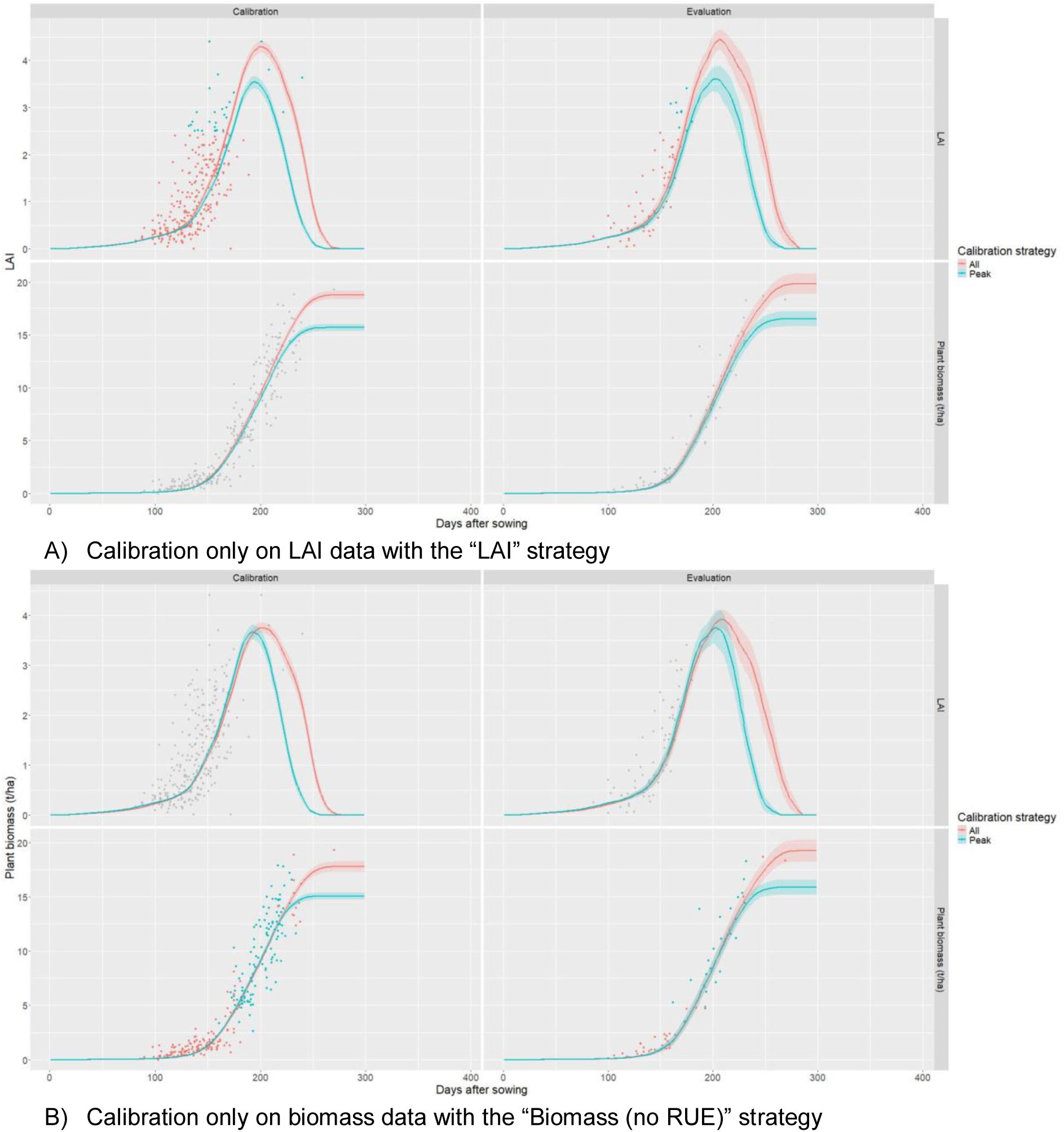
Simulated dynamics (continuous lines) with parameters estimated with two strategies for data availability (“All” in red, “Peak” in blue) and observations (individual points) for LAI and plant biomass variables when calibration is performed A) Only on LAI data with the “LAI” strategy and B) Only on biomass data with the “Biomass (no RUE)” strategy.

**Table 3.** Mean NRMSE values computed on entire or peak data, with two calibration options for data availability for LAI and plant biomass. Note that NRMSE values computed on the entire data with the calibration option “All” correspond to the ones obtained in Table 2 (*).

|  |  | LAI |  | Plant biomass |  |
| --- | --- | --- | --- | --- | --- |
| NRMSE computation | Calibration option | Calibration with “LAI” strategy | Evaluation | Calibration with “Biomass (no RUE)” strategy | Evaluation |
| All | Including all data | **0.48\*** | **0.50\*** | 0.23* | 0.24* |
|  | Including only peak data | 0.49 | 0.52 | **0.22** | **0.20** |
| Peak | Including all data | 0.53 | NA | 0.17 | 0.20 |
|  | Including only peak data | **0.45** | NA | **0.16** | 0.20 |

Overall, when aiming for the best simulations of plant dynamics over the whole season, the peak time option is more relevant for plant biomass simulation (as already pointed out with synthetic data), while the “All” option gives a better prediction for LAI simulation. This will be discussed in the following section.

## 4. Discussion

In this study we chose to focus on exploring the inclusion of different steps in the calibration process proposed and tested on synthetic data in Wallach et al. (2024). This calibration protocol was tested for the first time on experimental observed data for the STICS crop model with the objective of calibrating a new cultivar. Due to this specific objective the choice of parameters to estimate slightly differed (see section 4.1.). This study is also complementary to the ongoing AgMIP exercises since their aim is the intercomparison of various crop models whereas here we could focus on exploring the protocol for a single model and within a less contrasted range of situations. However, besides the choice of calibration steps, the choice of parameters to estimate and of target observed data (nature and timing) are two other major aspects.

### 4.1. Choice of plant parameters to include in calibration

The choice of model parameters to estimate is a key decision made by the modeler. This choice is firstly driven by data availability but since crop models like STICS have a high number of parameters, a selection has to be made. A consensus is to calibrate the most sensitive parameters, previously identified either by conducting a sensitivity analysis and/or by expert knowledge. However, to calibrate new cultivars as in this study, the varietal specificity of model parameters should also be considered in priority. Indeed, in STICS nearly half parameters are considered as varietal while the others are specific; this suggests that only varietal parameters need to be calibrated to parameterize a new cultivar. The decision process for parameters to calibrate is displayed as a decision tree in Figure S6 and detailed in Table S2. We therefore proceeded as follows: among the many parameters to be estimated, priority was given to those to which the model’s variables of interest are sensitive (Ruget et al., 2002). Parameters that are considered to be insensitive or not very sensitive are left at their default value; these are in particular the parameters involved in the model’s “satellite” calculations (i.e. those that do not have a direct impact on the functioning of the system in terms of growth and development). Among the sensitive parameters, the value of some is easily accessible through bibliography, measurements or expertise. Others are difficult to determine experimentally and must be estimated in some other way. If these parameters control a process that is highly variable in the dataset (e.g. a parameter that affects the effects of water stress, as the dataset contains situations ranging from low to high stress), then their value is estimated using a calibration algorithm. If this is not the case, these parameters are left at their default value (Table S2). Moreover, two very sensitive parameters were not calibrated (*extin* and *nbjgrain*), as the calibration algorithm could have used them to easily compensate for the estimation error of the main variables of interest, since the variables directly affected by these parameters were not measured in the dataset. We thus chose to calibrate 8 out of 52 with 4 for phenology (step 1 of the calibration procedure, see Figure 1), 2 for leaves (step 2), 2 for yield formation (step 5) that are routinely calibrated in STICS.

However the varietal character of STICS plant parameters can be questioned and is actually regularly updated with knowledge gain on the varietal specificity of some parameters, as is the case for radiation-use efficiency (*efcroiveg*, *efcroirepro*), notably because some recent breeding efforts targeted this trait (Govind et al., 2023; Stella et al., 2023). Indeed, for the calibration of modern cultivars like Rubisko, such a varietal specificity might be at play so we chose to add these parameters. Besides, in AgMIP exercises, some non-varietal parameters are routinely included because they are known to be sensitive and their varietal aspect could also be questioned so we chose to include *cgrain* and *irmax* parameters for grain number and yield prediction. Here we chose to keep a minimal set of parameters to estimate, most of them are indeed major parameters (only 2 submitted to selection during calibration), which could be a limitation of calibration performance, however the other pendant is the risk for over parameterization (Wallach et al., 2018).

Besides, the choice of initial parameter values can also impact calibration. Throughout this study we chose to set these initial parameters at the values of cultivar Arminda, similar to Rubisko in terms of precocity.

### 4.2. Choice of calibration/ evaluation method

Another major choice for calibration is the observed data to include in the calibration and evaluation steps. Indeed, the structure of the dataset can greatly impact parameter estimation (Wallach et al., 2021a). It is a standard practice to divide the observed dataset into a training subset for calibration and an evaluation subset to test the predictive ability of the estimated parameters. Similarly to what is done in the AgMIP calibration exercises (Seidel et al., 2018), we chose to manually divide the dataset by trying to minimize the dependence level between the two subsets, i.e. with different years and sites, but independence could not be fully achieved due to the near-complete design of the trials. In addition, Liu et al. (2018) estimated from a simulation study that calibration datasets may include at least 50% of available data, a condition fulfilled in our design; they also identified that the number of needed situations to properly calibrate and evaluate the model should be greater than five-fold the number of parameters to estimate. Our study also fulfilled this condition, with 67 situations to parameterize 10 to 12 parameters, close to the median of modelers’ practices (Seidel et al., 2018). However, more robust results could be achieved with a leave-one-out (LOO) method; LOO results in the lowest prediction error (Hernández-Ochoa et al., 2024; Wallach et al., 2021b), but may lead to overfitting when randomly applied, as some situations may be used many times in both calibration or evaluation subsets. Seidel et al. (2018) underlined that modelers often underestimated prediction errors with the frequent overlap of experimental sites between calibration and evaluation steps. To avoid it, Casadebaig et al. (2020) proposed a split of the data set into five parts to perform a cross validation loop without repetition of data for the evaluation step on each of the five parts successively. Finally, Hernandez-Ochoa et al. (2024) examined the best way to combine seasons and years for calibration and evaluation dataset; while phenological parameters were parameterized identically regardless of the combination, this was not the case for biomass and yield data, suggesting that LOO validation would benefit from some flexibility in the choice of years devoted to calibration or evaluation depending on the database. It suggests that the choice of a calibration and evaluation method depends on the type of parameter, but also that it should be specifically considered further in greater depth.

### 4.3. Calibration performance is limited by the structure of observed data rather than calibration steps

#### 4.3.1. Simulating dynamic variables enhances the scope of the model

The focus of this calibration study was to assess the impact of adding steps in the calibration process by including LAI and/or biomass data. Naively, one might assume that adding information during calibration will improve the goodness-of-fit. However, as Guillaume et al. (2011) previously pointed out, end-of-season variables such as yield are better predicted when dynamic variables are not included in the calibration process. Our findings confirm this statement as we found lower NRMSE values for grain yield and grain number at harvest when the calibration strategy excludes LAI and biomass data. Nonetheless, even if yield predictions are a major output of crop models, some dynamic variables are crucial to better understand crop response and widen their range of application. For instance, in the face of climate change, crop water consumption is a major stake strongly related to LAI dynamics: in that case, realistically simulated LAI dynamics are decisive (Hoek Van Dijke et al., 2020). Besides, assessing calibration performance on end-of-season variables only might be biased as compensation can occur by adjusting parameters related to yield; this is visible in our findings with higher values for parameter *vitircarb* for strategies resulting in lower biomass dynamics.

#### 4.3.2. Calibration strategy success depends on temporal distribution of observed data and chosen metrics

When comparing different calibration strategies on LAI and plant biomass, we found that simulated dynamics differ substantially. LAI dynamics differed for maximum peak value as well as senescence whereas biomass dynamics were very similar until 250 days after sowing before strongly diverging afterwards. This shows that the choice of calibration strategy, i.e. the choice of including LAI and/or biomass steps, had a major impact on the outputs. We found that LAI dynamics are better simulated when LAI data is integrated in the calibration process, with a better visual fit and lower NRMSE values. Biomass results are not as straightforward; end-of-season biomass was visually underestimated for most strategies but with no correlation with NRMSE mean values. This may result from an important lack of biomass points near maturity. Indeed when looking at individual fits per simulation unit, it appears that biomass NRMSE is mostly dependent from early growth up to flowering when lots of observations are available; on the contrary, the results of the calibration strategies begin to diverge at a later stage of growth when only few observations are available. There is also an effect of the subset division for evaluation; some strategies with satisfying visual fit for the training dataset tend to overestimate biomass for the evaluation dataset. Such a result indicates the evaluation step should be also deepened further, according to the available dataset.

#### 4.3.3. Recommendations for calibration

The initial results enabled the selection of the “LAI” and “Biomass (no RUE)” strategies as the most relevant options. However, a significant bias in the “LAI” strategy with regard to cereal yield was highlighted. Based on these results for this specific dataset, we would advise to use the strategy “Biomass (no RUE)” which relies on calibration of LAI parameters (maximum rate of gross daily increase of LAI *dlaimaxbrut* and maximum leaf lifespan *durvieF*) with using plant biomass measurements. Indeed, this strategy estimates realistic parameters’ values, and even if simulated biomass is not as visually satisfying as the “LAI” strategy, it is close (with NRMSE = 0.23 for this strategy vs. 0.22 for the “LAI” strategy on calibration dataset, and NRMSE=0.24 for both strategies on evaluation dataset). The results also pointed out that when including LAI and biomass parameters (i.e. RUE) in the same calibration step on biomass data, the algorithm preferentially adjusts RUE as it is more directly impacting plant biomass. Hence, we advise not to include RUE parameters when calibrating only on biomass data. This may also explain why surprisingly, the “LAI + Biomass” strategy is not the best one. When calibrating only RUE parameters on plant biomass (step 3 of the “LAI+Biomass” strategy, LAI parameters calibrated on LAI data on step 2, see Fig. 1) there is no overestimation of RUE parameters as for the “Biomass (with RUE)” strategy, there is less bias on LAI and biomass with lower MSE values but they are higher for grain number and grain yield. Besides, even if LAI is visually well simulated, biomass appears strongly underestimated with this strategy. As previously discussed, this biomass underestimation is due to the structure of the dataset in terms of temporal distribution of observations, so in other conditions calibrating on both LAI and biomass data might be a good strategy.

### 4.4. Insights from synthetic data on calibration procedures and their limits for real datasets

When testing different strategies for the calibration of Rubisko in STICS, we found that the structure of the observed dataset is limiting. There is notably a lack of LAI data at its peak maximum value and biomass data near maturity. We performed an exploration of data availability on synthetic LAI and biomass data in an attempt to distinguish crucial phases for the acquisition of experimental data. The synthetic data was built by sampling 5 points in noised simulated dynamics during a given period of time. We found very little differences in simulations obtained with parameters estimated on different periods of LAI/biomass data before, during, or after maximum LAI. Even though errors were overall very low, they were lower for the “Peak” strategy with minimum bias, indicating that performing calibration only on observed data during the LAI peak is the best option.

However, when going back to the actual observed data and applying this strategy, calibration was not improved, on the contrary LAI and biomass were noticeably underestimated. This shows that the application of calibration results obtained on synthetic data to a framework with real observed data could be limited. This can be explained by the very nature of synthetic data; since synthetic data is completely explainable by the model calibration can be facilitated. The different calibration results between synthetic and observed data could also be explained by the difference in data structure; indeed in the synthetic dataset each environment is equally represented with 5 samples distributed at regular time intervals whereas the observed dataset is more heterogeneous, and the total number of points is also much higher for the synthetic dataset with 335 points for both LAI and biomass vs 147 for observed LAI and 143 for observed biomass over the 67 calibration environments. Even though there is more data in the synthetic dataset, the acquisition of 5 LAI or biomass samples in a given environment is experimentally reasonable. Hence, even if exploring calibration on synthetic data can be useful, the results are not transferable to observed data for our case study. In any case, our results underline the need to acquire more data around the maximum LAI, which would allow a better simulation of plant biomass dynamics, and data throughout the whole season (including senescence and maturity) to avoid calibration strategies diverging due to parameter compensation issues.

### 4.5. Recommendations and perspectives

Based on the results obtained for this specific dataset, we would advise to use the strategy “Biomass (no RUE)”. We also recommend not to calibrate both LAI and RUE parameters on plant biomass data. We recommend caution when using synthetic data to explore calibration options as the obtained results are not systematically transferable to an observed dataset. Regarding experimental data acquisition, we advise for more balanced acquisition throughout growth for dynamic variables such as LAI or plant biomass, for a more balanced representation of the different phenological stages. For calibration purposes, LAI senescence should be measured as LAI instead of traditional senescence measures, i.e. percentage of senesced total surface so that the entire LAI dynamic can be described by the same variable.

As a perspective, model and calibration performance could be improved by increasing datasets in each situation; replacing LAI by fAPAR (fraction of photosynthetically active radiation) measured indirectly by remote sensing in models is a promising way to overcome the limitations of direct LAI measurement. This would notably enable more important and frequent data acquisition. This has already been successfully used with satellite fAPAR products, which helped to better characterize some critical parameters controlling the phenology of the ORCHIDEE process-based vegetation model (Bacour et al., 2015). Calibration performance could also be improved by modifying the weights of time points for the computation of RMSE, notably to account for heterogeneous data acquisition overtime; in our case study, that is to say adding more weight to end-of-season points for dynamic variables or less weight to begin-of-season points that are superfluous in terms of the information provided. Here this could be done by choosing weights inversely proportional to the number of observations in a given temporal window to balance the weights of observations throughout growth (He et al., 2024). More generally, adding weight to a critical variable and/or temporal window for RMSE computation could also be a choice made by the modeler to highlight the importance of a given process and/or objective such as prediction at maturity. Visual assessment of the goodness of fit of simulated variables is a valuable tool for model calibration, although it can sometimes conflict with quantitative metrics such as NRMSE, so the combination of these two approaches could offer a compelling solution. Machine learning methods such as convolutional neural networks (CNNs), which are widely used for tasks such as plant disease detection (Hasan et al., 2020) and medical image analysis (Cicero et al., 2017), could complement traditional validation techniques by serving as a proxy for human expertise. Using these AI methods to assess model fit is a promising way forward, potentially bridging the gap between quantitative metrics, and empirical visual assessments. However, such approaches face significant challenges, particularly in terms of the quality and quantity of training data required (Alzubaidi et al., 2021).

## Supplementary Material

**Table S1.**
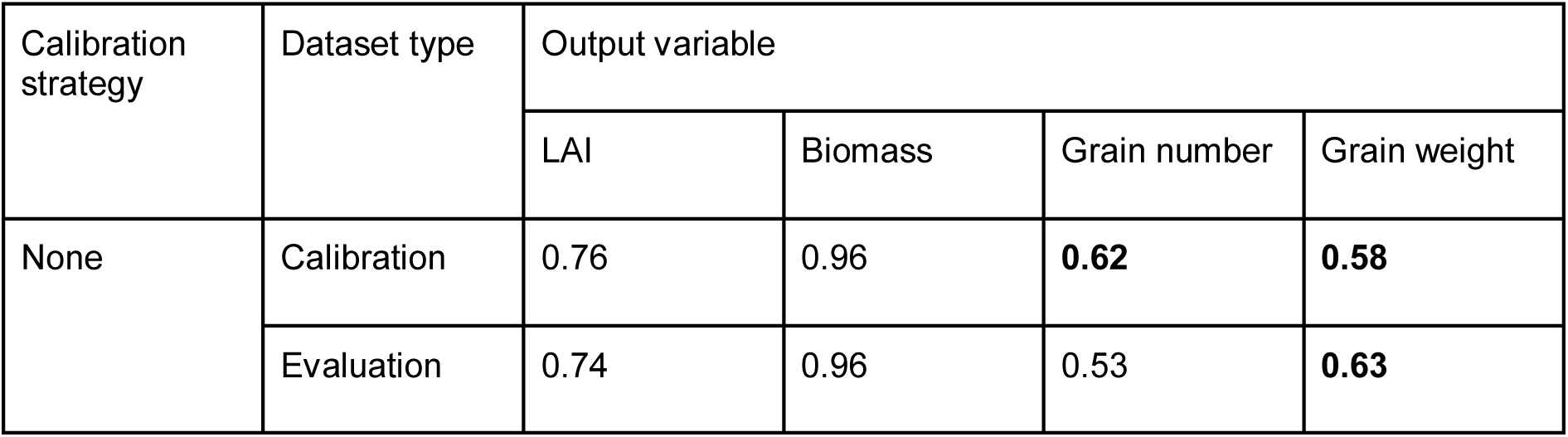
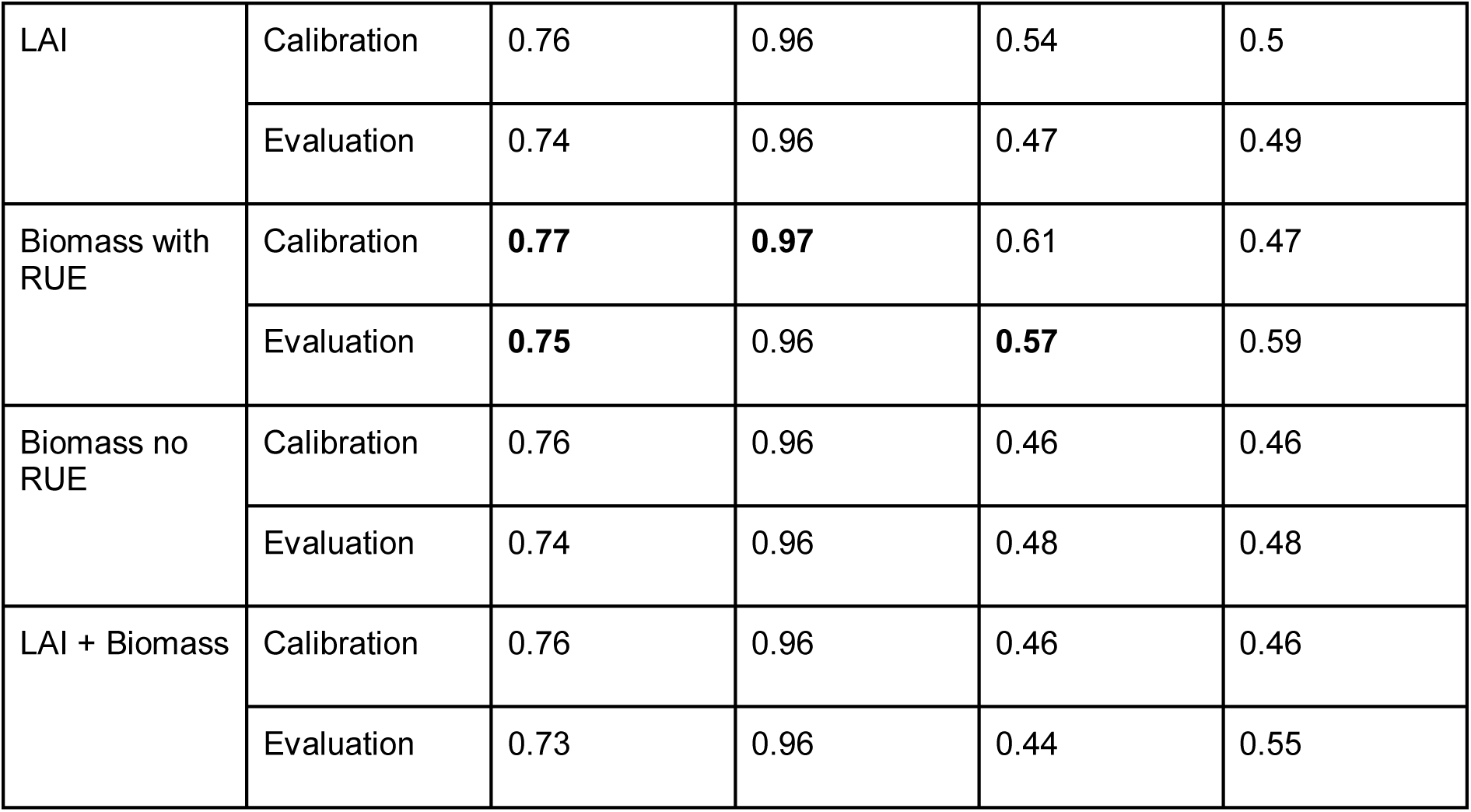
Pearson correlation coefficients between observations and simulations for the calibration and evaluation datasets for each calibration option.

Table S2 - Varietal parameters of the STICS crop model (see xlsx file)

**Figure S1.**
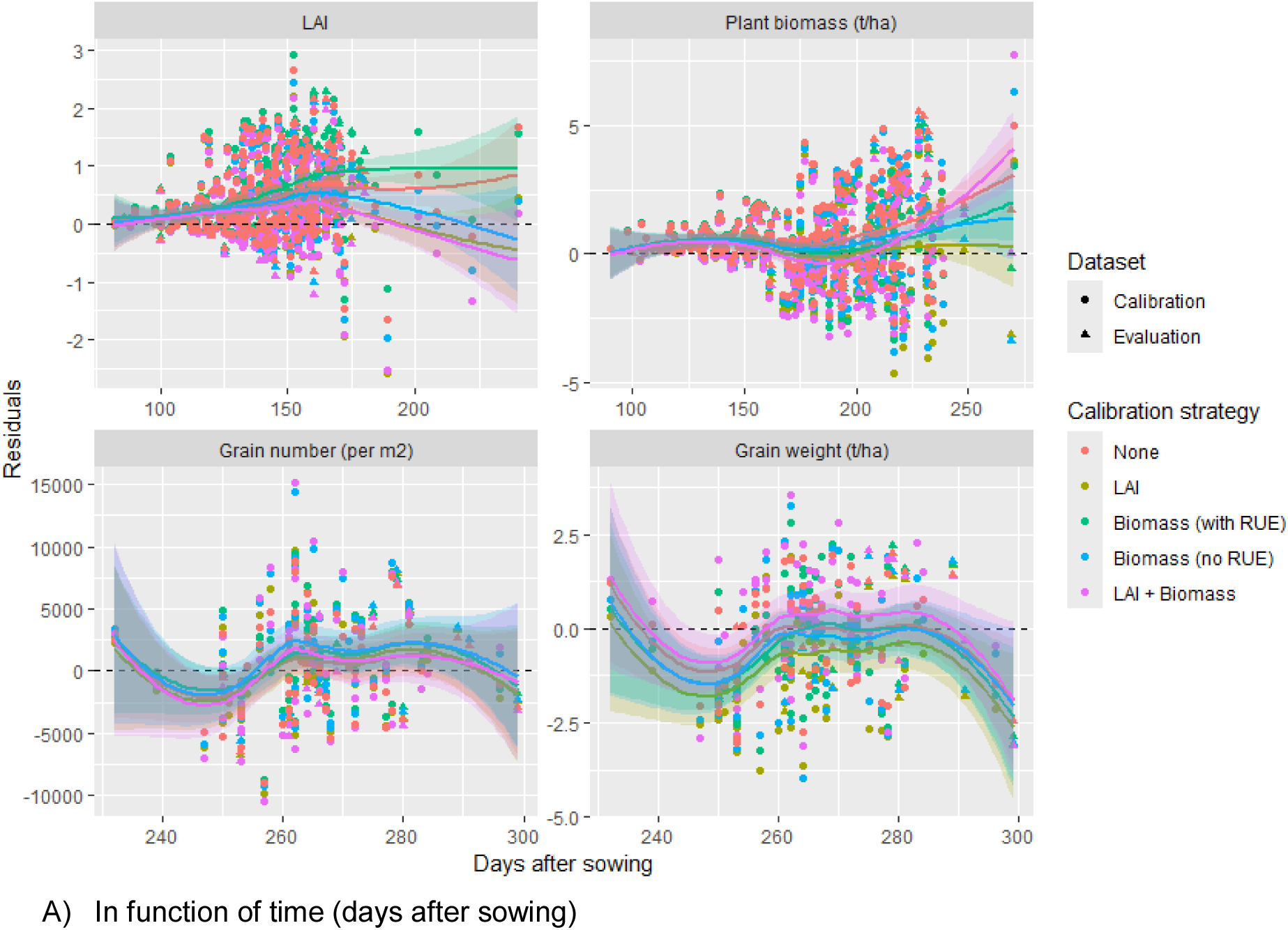
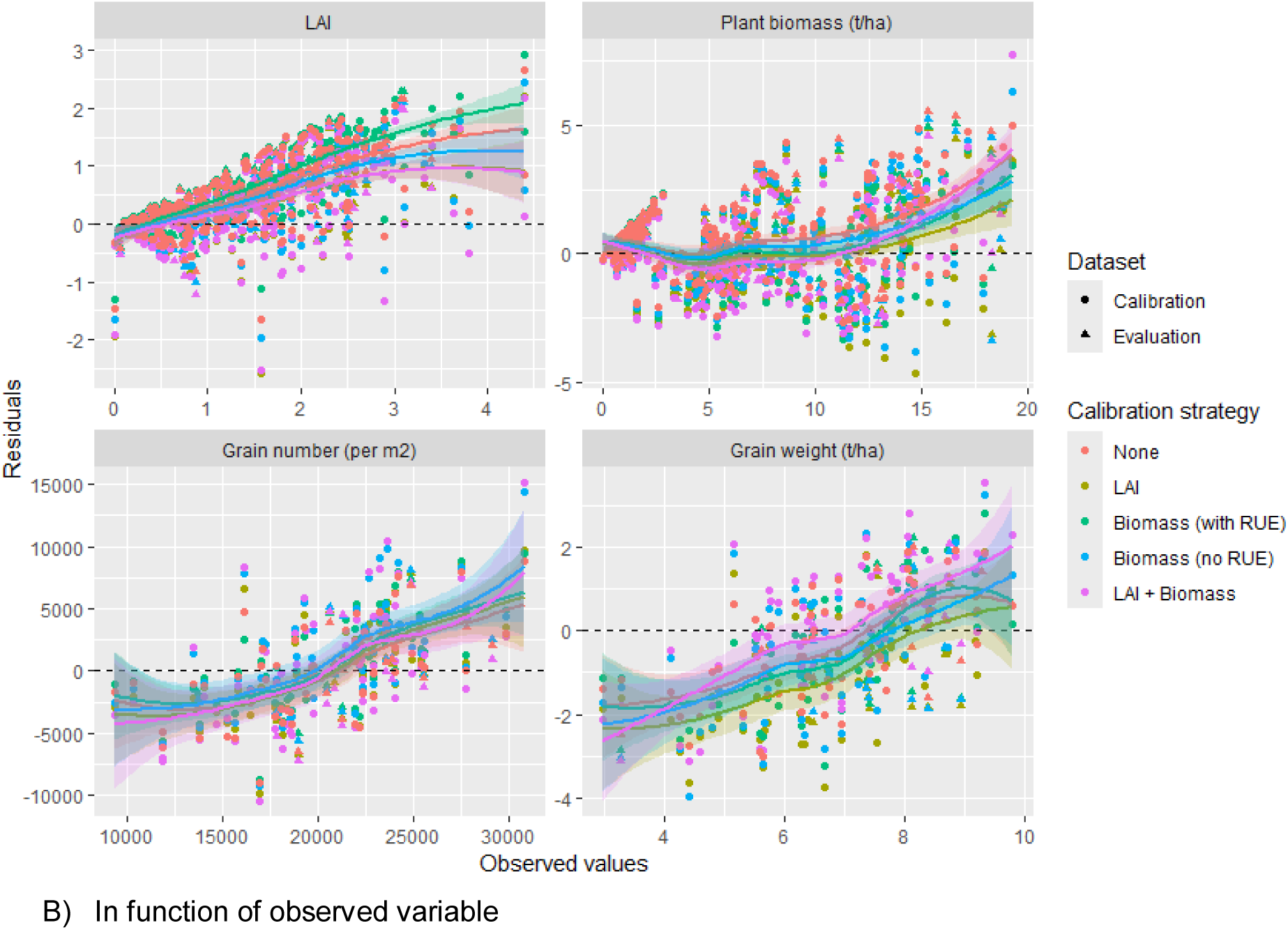
Residuals between observations and simulations in function of A) Time (days after sowing) or B) Observed variable for LAI, plant biomass (t/ha), grain number (per m^2^) and grain weight (t/ha). Point shape correspond to the dataset type (calibration or evaluation), colors correspond to the different calibration strategies and lines correspond to Loess non-parametric regression.

**Figure S2.**
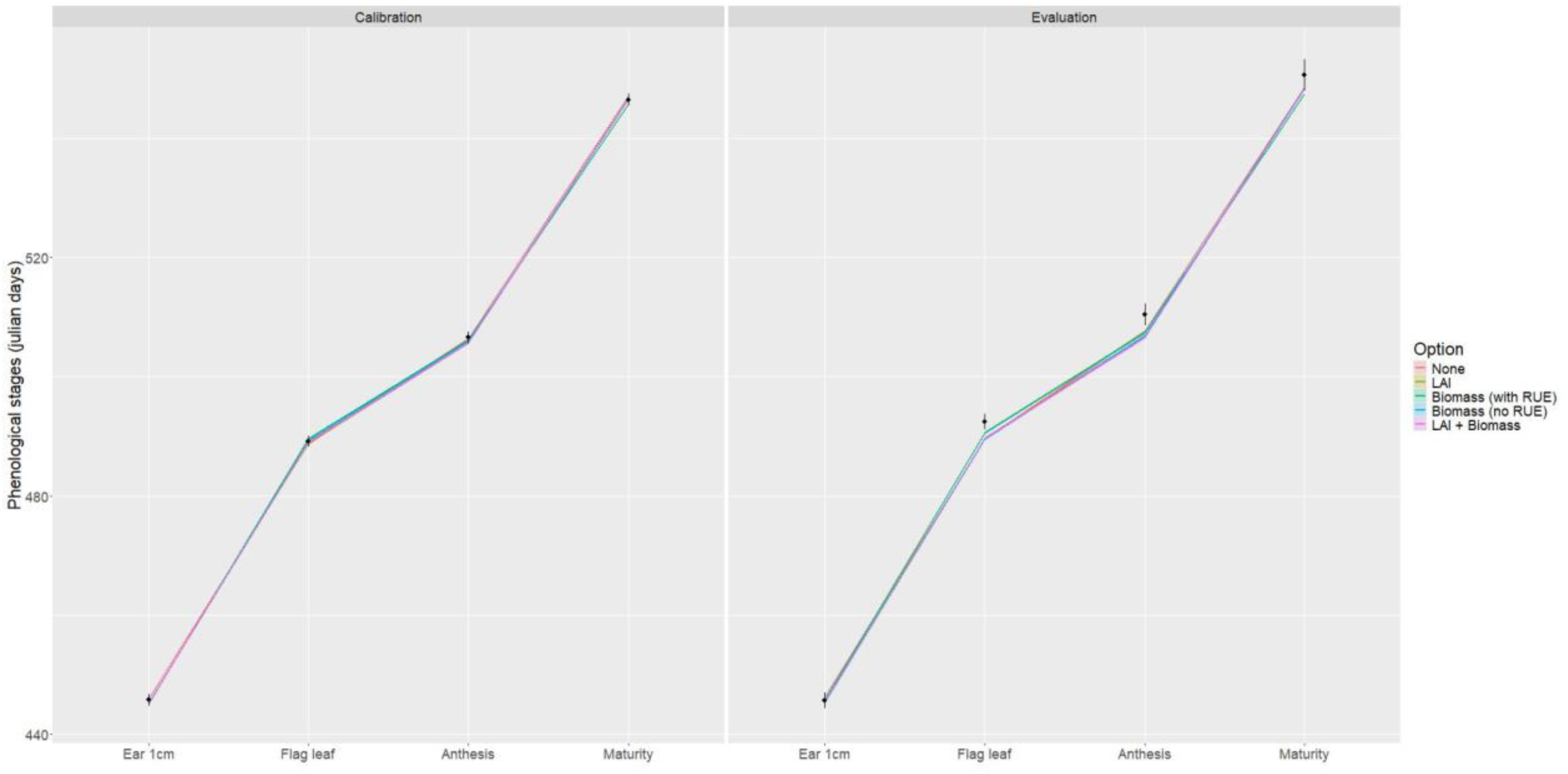
Simulated versus observed phenological stages ear 1cm, flag leaf, anthesis and harvest. Colored lines represent simulated mean phenological stages for the different calibration strategies with a ribbon for standard error around the mean. Black points represent observed mean phenological stages and the error bars represent the standard errors around the mean.

**Figure S3.**
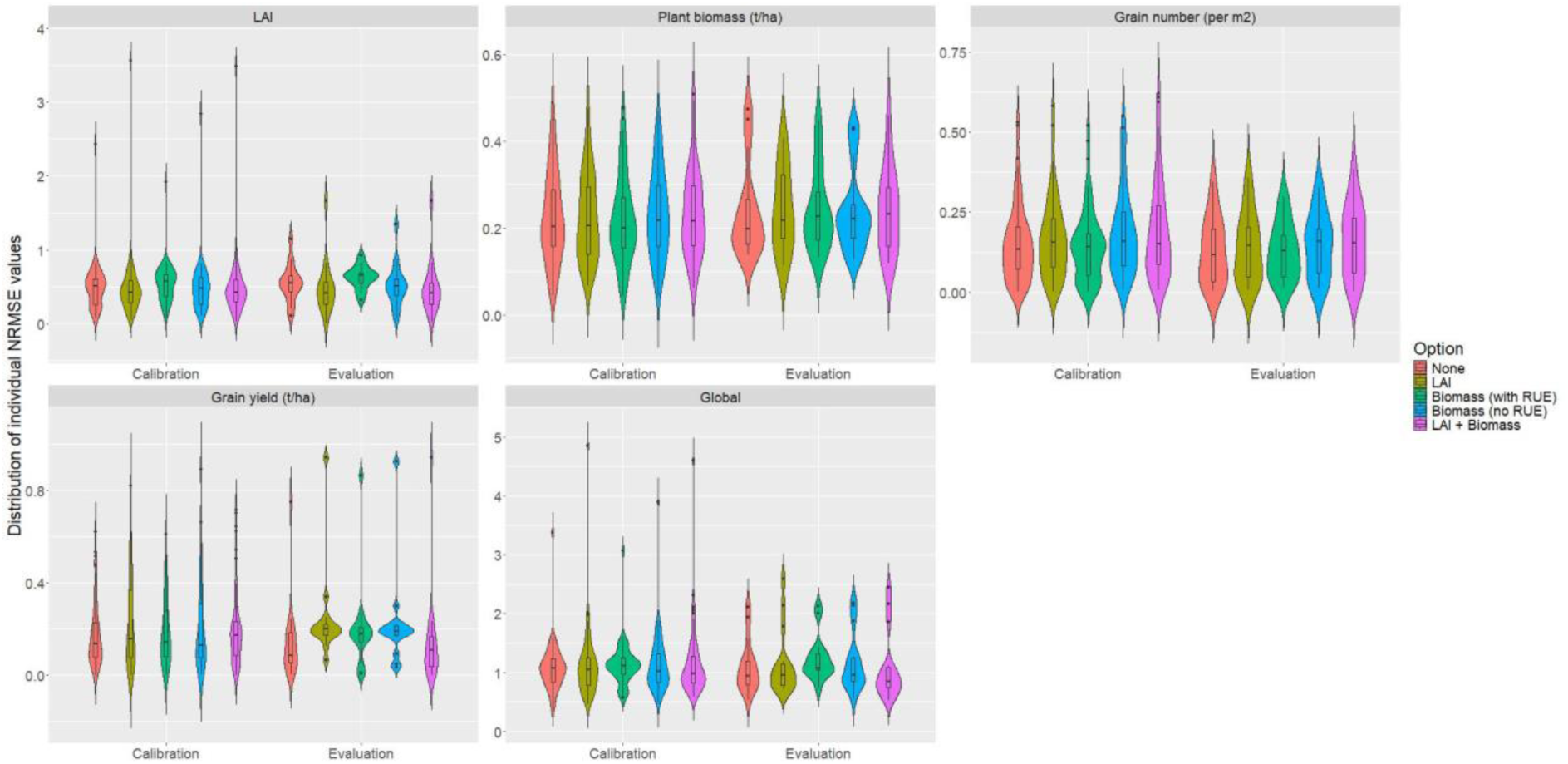
Distribution of individual NRMSE values faceted for different target variables (LAI, plant biomass, grain number, grain yield and global), coloured for different calibration strategies (None, LAI, biomass with RUE, biomass no RUE and LAI + biomass) and for both the calibration and evaluation datasets. The scale of the errors for each variable is different.

**Figure S4.**
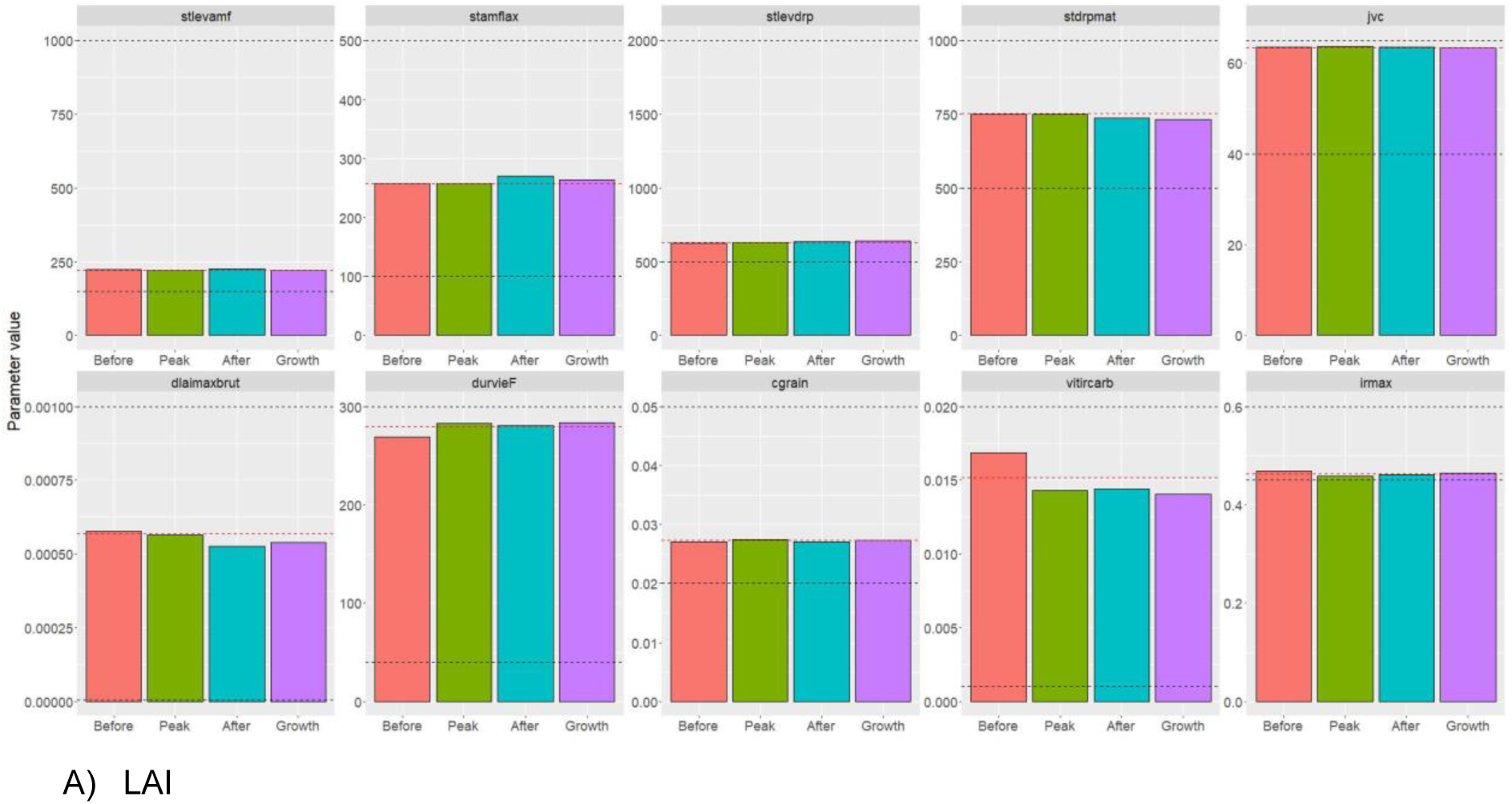
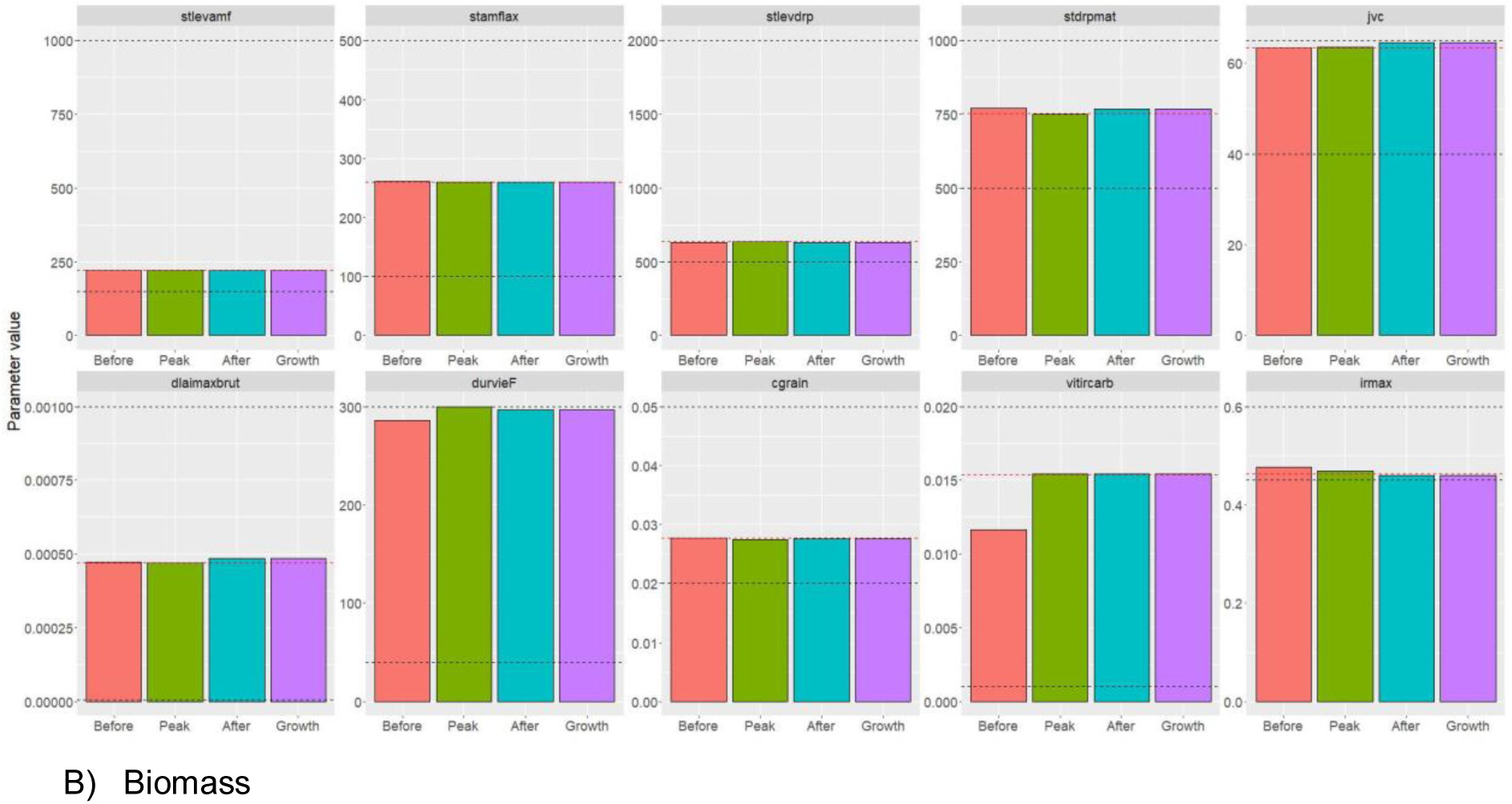
Estimated values of calibrated parameters for each calibration strategy for synthetic data availability before, during, after the LAI peak and throughout growth for calibration on A) LAI and B) Biomass. Black dashed lines represent lower and upper bounds, red dashed lines represent estimated values found on actual observed data.

**Figure S5.**
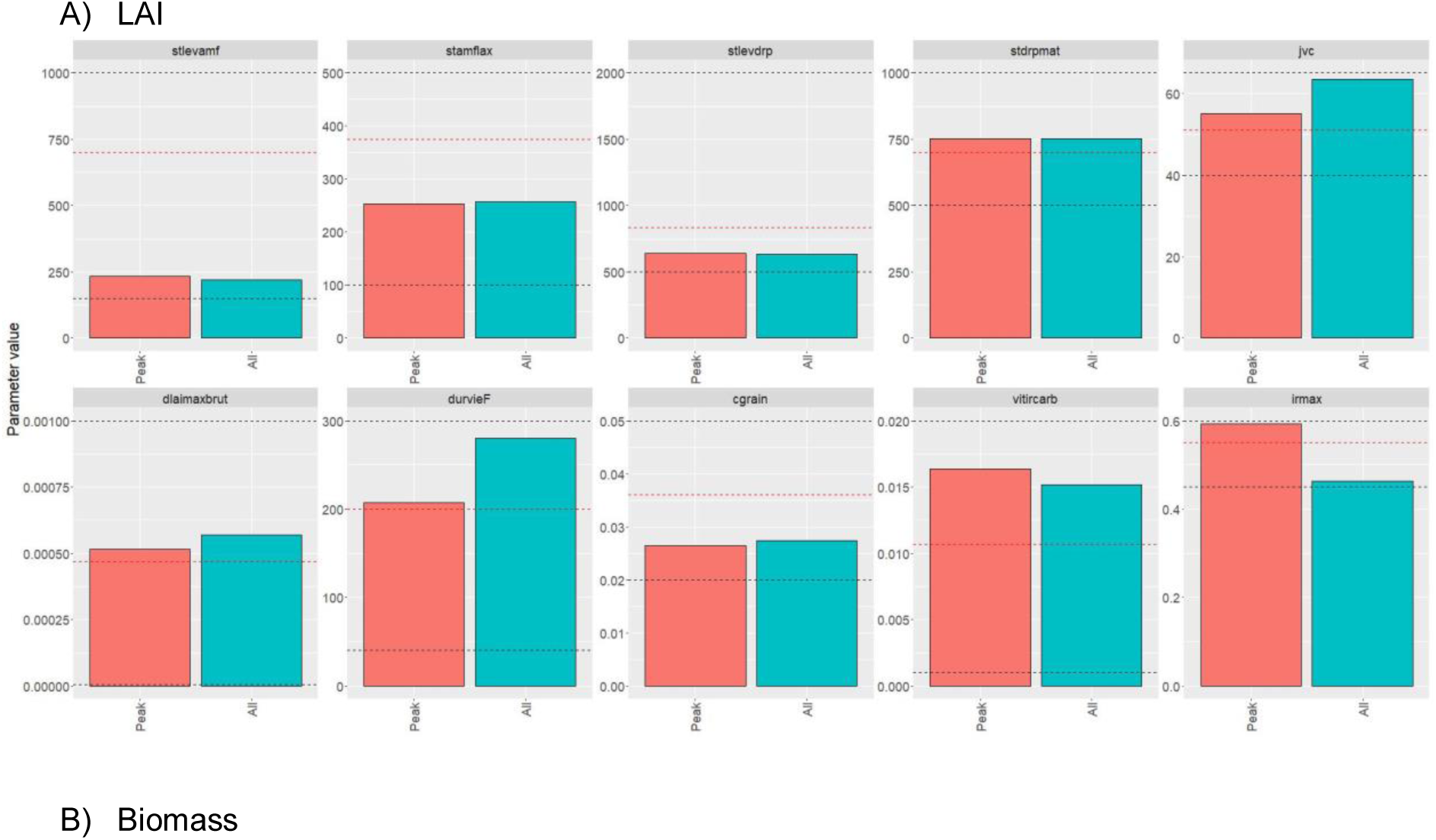
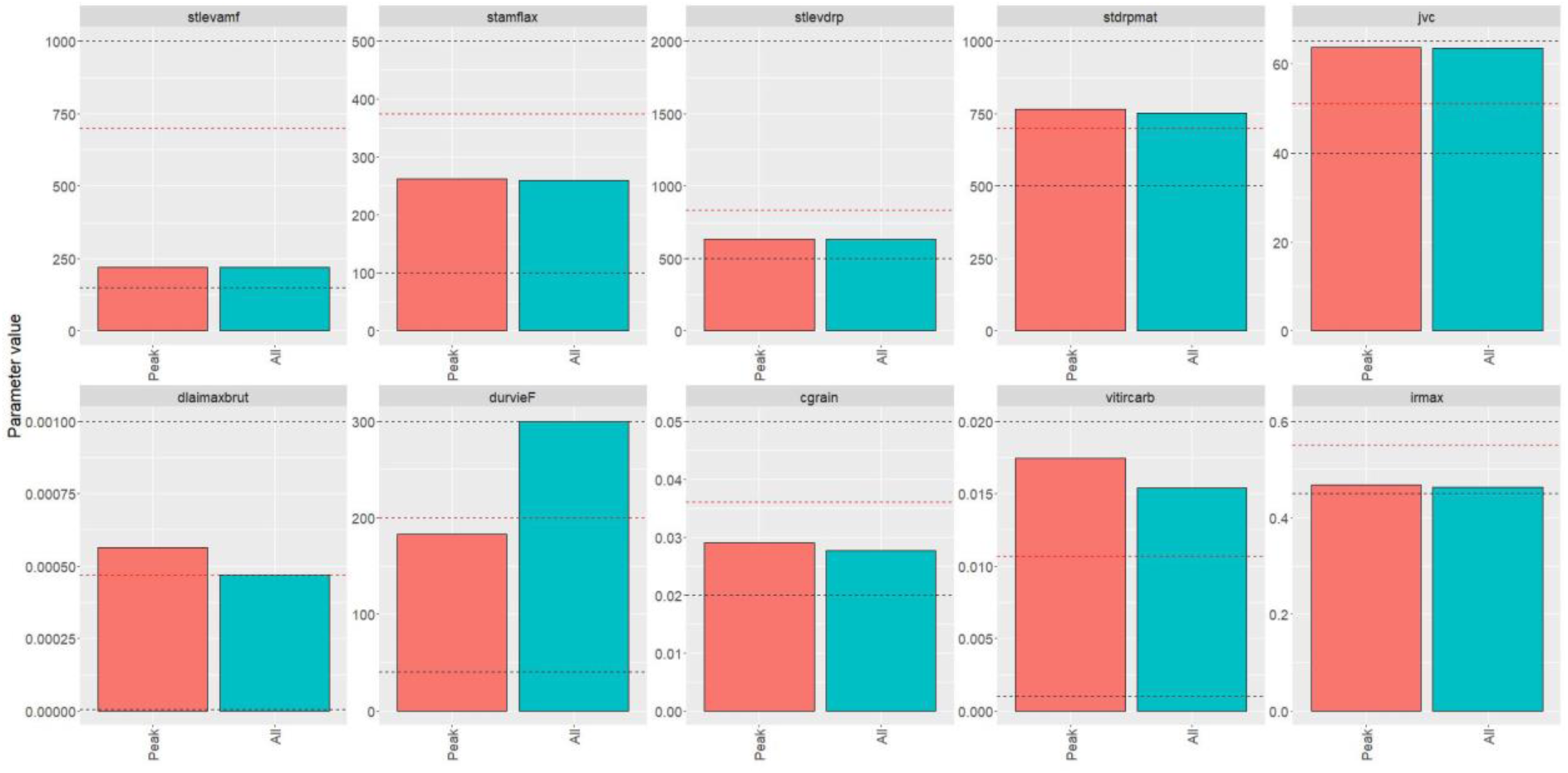
Estimated values of calibrated parameters for each calibration strategy of data availability during the LAI peak and throughout growth for calibration on A) LAI and B) Biomass. Black dashed lines represent lower and upper bounds, red dashed lines represent initial parameter values.

**Figure S6.**
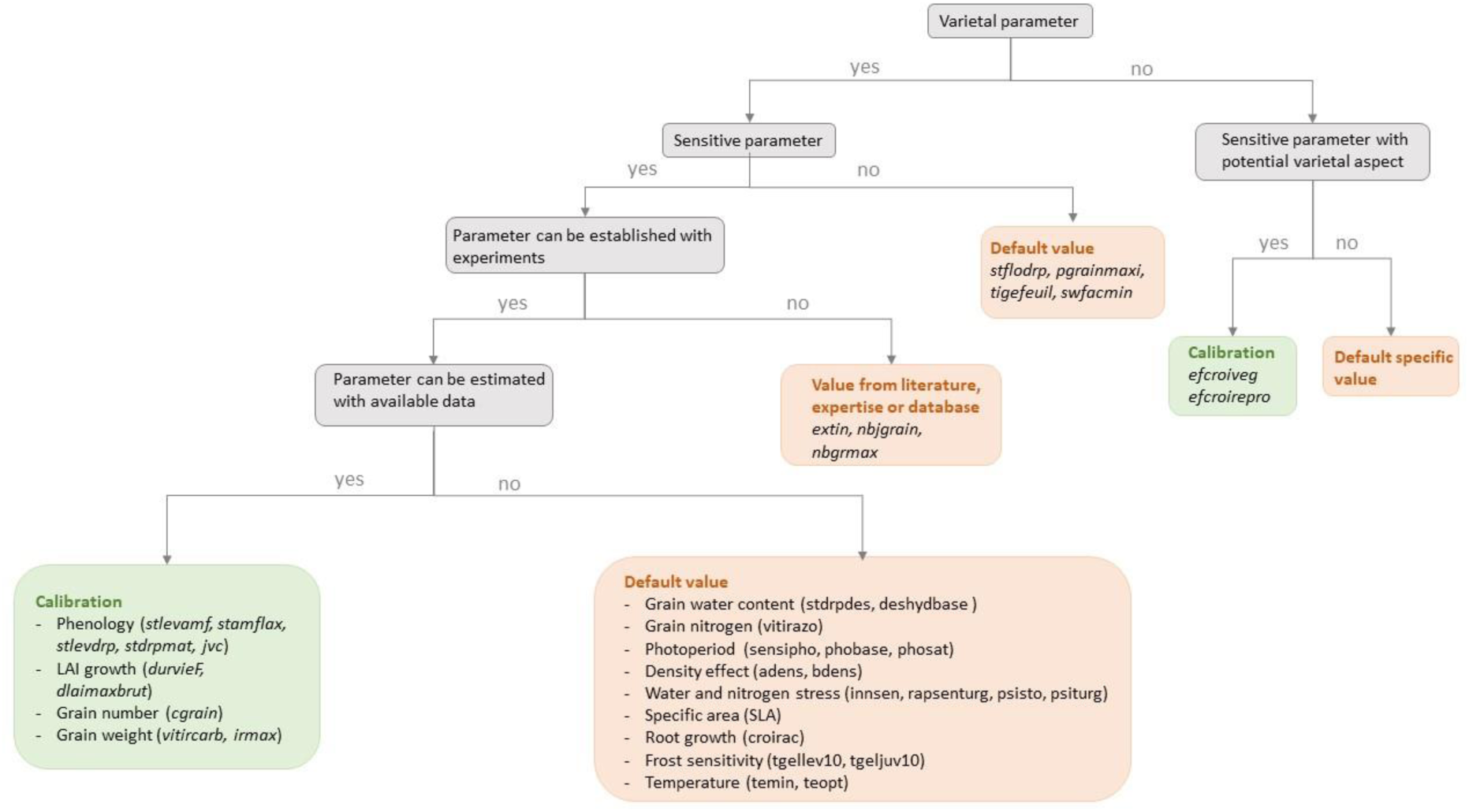
Decision tree for the selection of parameters to integrate in the calibration process for winter wheat in the STICS crop model

